# DNA flexibility regulates transcription factor binding to nucleosomes

**DOI:** 10.1101/2024.09.02.610559

**Authors:** Luca Mariani, Xiao Liu, Kwangwoon Lee, Stephen S. Gisselbrecht, Philip A. Cole, Martha L. Bulyk

**Author notes:** Correspondence should be addressed to M.L.B. and L.M.

## Abstract

Cell fate decisions are controlled by sequence-specific transcription factors (TFs), referred to as ‘pioneer’ factors, that bind their target sites within nucleosomes (’pioneer binding’) and thus initiate chromatin opening. However, pioneers bind just a minority of their recognition sequences present in the genome, suggesting that local sequence context features may regulate pioneer binding. Here, we developed PIONEAR-seq, a highly parallel sequencing-based biochemical assay for high-throughput analysis of TF binding to nucleosomes on nucleosome positioning sequences. Using PIONEAR-seq, we characterized the pioneer binding of 7 human pioneer TFs. Comparison of TF binding to nucleosomes based on the synthetic Widom 601 (W601) model sequence versus three different genomic sequences revealed that the positional preferences of these TFs’ binding to nucleosomes (*i.e.*, dyad, periodic and end binding) is determined by the broader sequence context of the nucleosome, rather than being a property intrinsic to the TF. We propose a model where the flexibility and rigidity within nucleosomal DNA regulate where pioneers bind within nucleosomes. Our results suggest that the broader physical properties of nucleosomal DNA represent another layer of *cis*-regulatory information read out by TFs in eukaryotic genomes.

## MAIN

In eukaryotes, gene expression programs are dynamically regulated by changes in the accessibility of chromatin for transcription factor (TF) binding to specific DNA motifs (6-10 bp). TFs that can pioneer inaccessible chromatin for subsequent transcriptional activity thus serve as gatekeepers of cellular differentiation^1,2^. However, it remains unclear which TFs are able to target sites within nucleosomes and what controls their pioneer binding.

Several, not mutually exclusive models for how TFs access their nucleosomal DNA binding sites have been proposed^3^, such as competition between TFs and histones for DNA binding^1,4,5^ or spontaneous, partial unwrapping of nucleosomes providing transient access to DNA target sites^6,7^. Pioneers have been found to exhibit differences in their preferential engagement with certain regions of nucleosomes, in particular at: the dyad^8^, where the single DNA gyre is anticipated to be more accessible than elsewhere within the nucleosome where there are two DNA gyres^9^; the ends^10–12^, where the DNA may spontaneously unwrap from the histone octamer, thus, providing transient access to DNA target sites^6,7^; internal positions of nucleosomes through DNA distortion^13^; or positions spaced with approximately 10-bp periodicity^14^, where TF binding sites in the major groove of the DNA face outward^9^. While some TFs may target different recognition sequences on nucleosomes than on free DNA^10,12,15–17^, such differences do not explain differences among TFs in their positional preferences of binding to nucleosomes.

Numerous biochemical studies of TF binding to nucleosomes have used nucleosomes assembled on synthetic DNA sequences selected for high affinity for the histone octamer^14^, in particular those based on the Widom 601 (W601) template^10–13,18–20^. A recent large-scale *in vitro* selection (“NCAP-SELEX”) survey of extended DNA binding domains of 220 human TFs classified them according to their positional preferences of binding to nucleosomes built on completely randomized DNA sequence^14^. However, it remains unclear whether results from binding to purely synthetic nucleosomal DNAs generalize to binding to nucleosomes assembled on genomic DNA sequences. For example, nucleosome N1 within the ALB enhancer (*ALBN1*), which is targeted *in vivo* by the pioneer factor FOXA during liver development^21^, exhibits FOX-specific sites in close proximity (<15 bp) to the dyad^8,22^; in contrast, results from the NCAP-SELEX survey found FOX TFs to be end and periodic binders^14^. As another example, the CX3 chemokine receptor 1 (*CX3CR1*) nucleosome, which is pioneered by PU.1 and CEBPA TFs to promote macrophage identity, contains their binding motifs near the expected dyad and is bound by them when assembled *in vitro* into either mononucleosomes^23^ or compacted nucleosome arrays^24^. In contrast, CEBPB, a close paralog of CEBPA, exhibited strong end-binding preference in the NCAP-SELEX survey^14^. As yet another example, a nucleosome from the *LIN28B* distal enhancer, which is pioneered *in vivo* by SOX2 and OCT4 during reprogramming^25^, was bound by both SOX2 and OCT4 *in vitro*^16^, whereas no or weak binding by SOX2 or OCT4, respectively, was detected on W601-based nucleosomes carrying their binding motifs at multiple positions^10^. Moreover, the structures of OCT4 bound to W601 versus *LIN28B* nucleosomes differ considerably: OCT4 binds at the entry-exit sites on W601 nucleosomes using exclusively its POU DNA binding domain (DBD)^10^, while on *LIN28B* nucleosomes (as well on nucleosomes based on the n*MATN1* genomic locus) it binds using both the HD and POU DBDs on the linker DNA, which is spatially proximal to the dyad, and also using the activation domain, which interacts with the histone H3 and H4 N-terminal tails^26^.

A cryo-EM study of a nucleosome containing the *ALBN1* enhancer DNA sequence found that the histones were more weakly associated with that DNA than in a nucleosome containing W601 template, suggesting that different DNA sequences may have different accessibility to DNA binding proteins^27^. Synthetic DNA sequences preferentially bound by the histone octamer display 10-bp periodicity in single nucleotides and dinucleotides^14,18^, in agreement with that found in nucleosome positioning sequences in eukaryotic genomes^28^. These periodic oligonucleotide signals facilitate DNA sequence bending at precise positions around the histone octamer, thus inducing the formation of stable nucleosomes^29^. A single-molecule study found that the unwrapping of one DNA end of a W601-based nucleosome can stabilize the other end^30^, suggesting that differences in DNA composition across nucleosomes might result in differential access to the ends of nucleosomes. Furthermore, results from several *in vivo* studies indicate that pioneers bind *in vivo* to only a minority of the genomic sites encompassing their recognition motifs^31–33^. Altogether, these results raise the possibility that the DNA sequence context of nucleosomes may control the utilization of TF binding sites by pioneers.

To address this question, we developed PIONEAR-seq technology and used it to characterize binding by 7 human pioneer TFs to nucleosomes based on W601 versus three genomic sequences, each of which we tiled with random DNA segments. This approach provides a platform to interrogate the sequence context features that regulate TF binding to nucleosomes.

## Results

### PIONEAR-seq assay design

To determine the effects of nucleosome DNA composition on the sequence specificity and positional preferences of pioneer binding, we developed PIONEAR-seq (Pioneer Interactions On Nucleosomal Elements After Reconstitution followed by sequencing) for highly parallel analysis of the nucleosomal DNA binding preferences of TFs (**Fig. 1a**). Briefly, in PIONEAR-Seq, we tiled across 147-bp nucleosome positioning sequences (’templates’) every 3 bp with 20-bp random regions (N_20_ tiles) (’PIONEAR-seq design of the Input Library’ in **Supplementary Information**). Here, we used four well-characterized templates, three of which represent *in vivo* regulatory elements associated with three genomic nucleosomes (*ALBN1, CX3CR1, NRCAM*)^23,27^ and the high-affinity W601 sequence^34,35^ (**Supplementary Table 1)**. Merging the resulting four template sublibraries resulted in a library of ∼4 x 10^13^ unique DNA sequences (**Supplementary Table 2)**. The DNA libraries were dialyzed with histone octamers comprising the canonical human histones (H2A, H2B, H3.1 and H4) (“Histone octamer assembly and purification” in **Supplementary Information**, and **Extended Data Fig. 1a)** to form nucleosome core particles (“Nucleosome assembly” in **Supplementary Information**, **Extended Data Fig. 1b**). We confirmed correct nucleosome assembly by MNase-seq profiling (**Extended Data Fig. 1c**). Insertion of sequences containing known TF binding sites did not modify the overall accessibility profile of the nucleosome core particles (NCPs) assembled on single templates containing TF binding sequences at different positions (**Extended Data Fig. 1d, Supplementary Table 3**) nor on the template sublibraries (**Extended Data Fig. 1e,f).** The resulting ‘*NCP input library*’ was used in selections for binding by GST-tagged TFs in four rounds of nucleosome assembly, TF selection and PCR amplification (**Fig. 1a**). Selections using GST alone were performed as a negative control. After four rounds of selections, libraries were sequenced and analyzed for enrichment of TF DNA binding sequences as compared to the input libraries and for the positional preferences of these sequences within the nucleosomes.

**Figure 1:**
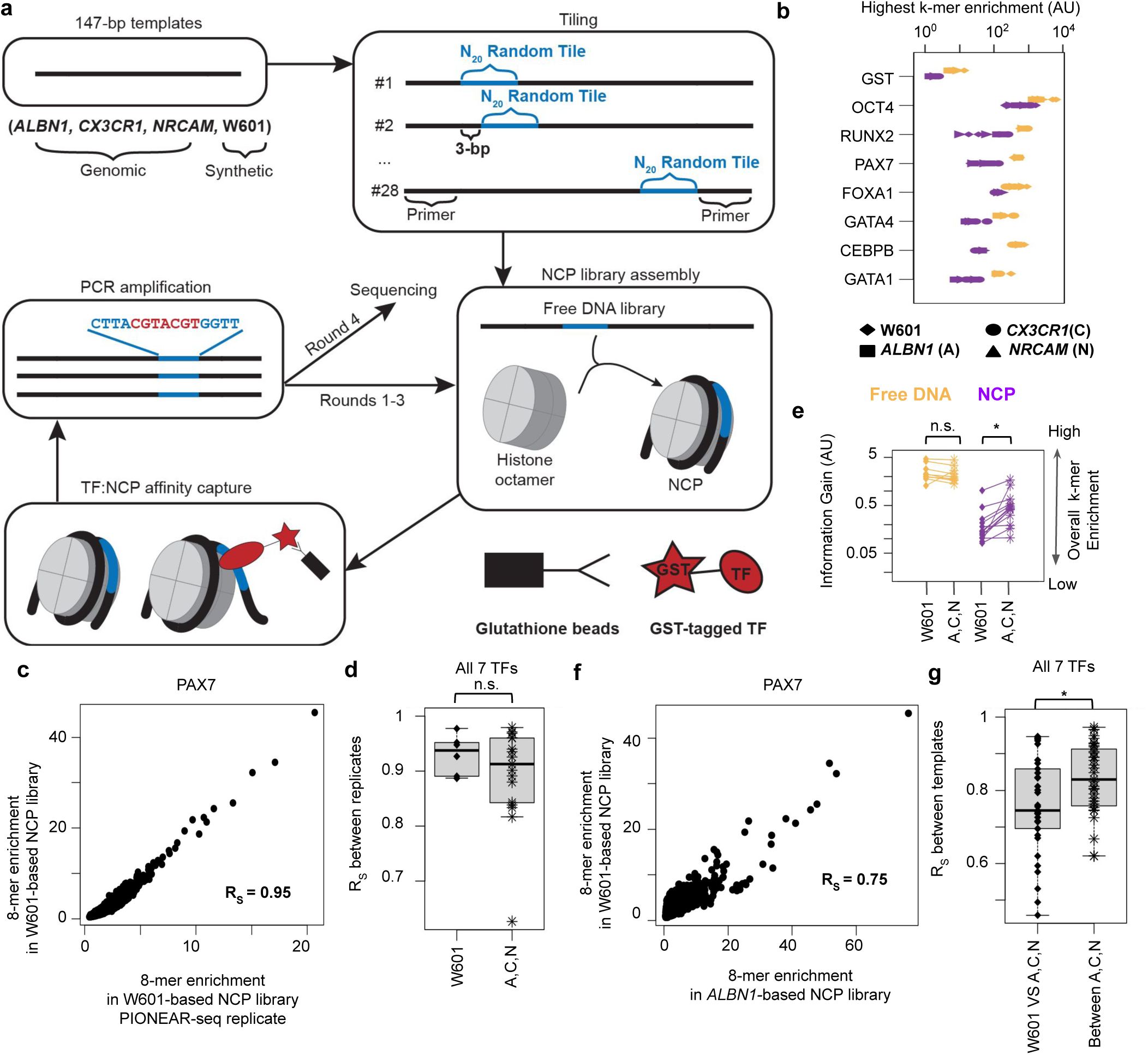
PIONEAR-seq assay reveals TFs with specific binding to nucleosomes. **(a)** PIONEAR-seq assay schema. DNA sequences that favor nucleosome positioning (’147-bp Templates’) are tiled with 20-bp random regions (N_20_ random tiles, blue) in the internal regions between fixed primers and pooled together to form the free DNA input library (’Tiling’), which is then assembled into NCPs to form the NCP input library using previously reconstituted histone octamers (’NCP Library Assembly’). NCPs are then incubated with GST-tagged TFs and pulled down using GST-specific beads (’TF:NCP affinity capture’). The selected DNA libraries are PCR-amplified, selected by 3 additional rounds, and then sequenced to determine TF binding sites within the N_20_ tiles (’PCR amplification’). **(b)** Comparison of the highest k-mer enrichment in libraries based on distinct templates (symbols), selected by the indicated TFs (or the GST mock control) and assembled on NCPs (purple) versus kept as free DNA (yellow). **(c)** Representative example of correlation of *k*-mer enrichment (Spearman R = 0.96) between NCP library replicates upon PIONEAR-seq selection. **(d)** For the 7 TFs in (b), the correlation of *k*-mer enrichment between PIONEAR-seq replicates in W601 libraries versus libraries based on the genomic templates (*ALBN1, CX3CR1* and *NRCAM*) (Wilcoxon Rank Sum Test, *P* = 0.53). **(e)** For the 7 TFs in (b), comparisons of information gain of libraries based on W601 versus the genomic templates and selected either as NCP (purple) or free DNA (yellow) (**Extended Data Fig. 2c)**. **(f)** Representative example of correlation of *k*-mer enrichment (Spearman R = 0.78) between NCP libraries based on W601 and the genomic templates upon PIONEAR-seq selection. **(g)** For the 7 TFs in (b), the correlation coefficients of *k*-mer enrichment between W601 and each genomic template are significantly lower than among the genomic templates (Wilcoxon Rank Sum Test, *P* = 9.8 x 10^-3^).

### Nucleosomal sequence context influences TF binding to nucleosomes

We assayed 11 full-length human TFs in duplicate by PIONEAR-seq (**Extended Data Fig. 2a, Supplementary Table 4)**: 10 that have been reported to act as pioneers either *in vitro* or *in vivo* (ASCL1^36^; CEBPB^37^; FOXA1^38^; GATA1^39^; GATA4^38^; KLF4^16^; OCT4/POU5F1^16^; PAX7^17^; PU.1/SPI1^37,40^; RUNX2^41^ and one, ZNF143, that has not been reported to bind nucleosomal DNA. Since pioneers typically serve as regulators of cell type specification, we hypothesized that ZNF143, a key developmental regulator expressed across cell types^42^, would not serve as a pioneer. In parallel, we assayed these TFs for binding to the free DNA input libraries.

To inspect the sequence specificity of TF binding in an unbiased manner, we analyzed the randomized tiling sequences computationally using a *de novo* motif discovery method^43,44^ which scores DNA sequences of length *k* (’*k*-mers’) for enrichment as compared to the input. For each value of *k,* we then combined the enrichment of multiple *k*-mers per library to calculate the information gained by the input library during selection, which can be quantified by the Kullback-Leibler divergence^44^, and thus inferred each TF’s optimal DNA binding site length in each template sublibrary^43^ (**Supplementary Information)**. In subsequent analyses we assigned to each TF the average optimal *k*-mer length among its libraries, which was 10-mers for OCT4; 9-mers for FOXA1, KLF4, PU.1 and ZNF143; and 8-mers for the remaining TFs. The TFs’ top enriched *k*-mers in free DNA libraries matched their known DNA binding specificities^45^ (**Extended Data Table 1, Supplementary Table 5**).

To evaluate TF DNA binding enrichment in each library, we used two scores: 1) the enrichment (**Fig. 1b, Extended Data Fig. 2b**) of a TF’s most preferential DNA binding site *k*-mer, and 2) the Kullback-Leibler divergence (*information gain*) (**Extended Data Fig. 2c,d**) for comprehensive scoring over all the *k*-mers of optimal length enriched in a library (as described previously for SELEX-Seq data^43,44^). *K*-mer enrichment was highly reproducible across library replicates (median Spearman correlation coefficient R = 0.93; **Fig. 1c**) and was indistinguishable on libraries based on W601 versus genomic templates (**Fig. 1d**). According to both scores, TFs consistently exhibited reduced binding specificity on NCPs than on free DNA (**Fig. 1b, Extended Data Fig. 2b-d),** consistent with prior reports that nucleosomes typically limit DNA binding^14,23^. Seven of the 11 TFs - CEBPB, FOXA1, GATA1/4, OCT4, PAX7 and RUNX2 - exhibited significant sequence-specific binding to NCP libraries as compared to GST negative controls (**Fig. 1b, Extended Data Fig. 2c**). These 7 TFs’ most enriched *k*-mers in binding to nucleosomes were almost identical to those from binding to free DNA (**Extended Data Table 1, Supplementary Table 5**), suggesting that other chromatin features may contribute to the preference for some TFs to recognize more degenerate sequence motifs on nucleosomes *in vivo*^15,16^. Of the remaining 4 TFs, ZNF143 has not been shown to bind nucleosomes, ASCL1 has been shown to require heterodimerization with a bHLH co-factor for detectable *in vitro* nucleosome binding^23^, nucleosome binding by KLF4 has been shown to be low or below detection in similar biochemical assays^23,16^, and PU.1 binding has been found to be restricted by nucleosomes and to bind only at the exit/entry sites of nucleosomes^19,46^ in a region beyond where we tiled with 20-bp random sequences. To further corroborate the results we obtained by PIONEAR-seq, we looked at the published E-MI diagonal profiles from lig200 NCAP-SELEX data^14^ for the same TFs or very close paralogs thereof (**Extended Data Fig. 2e,f)**. These profiles, which provide the positions of the TF DNA binding sites, revealed that TFs (or close paralogs) that bound nucleosomes in PIONEAR-seq assays exhibited E-MI profiles typical of nucleosome-binding TFs, which was not the case for TFs (or close paralogs) that did not bind nucleosomes in PIONEAR-seq (**Extended Data Fig. 2g,h)**.

We noticed that information gain was significantly lower on W601-based NCP libraries than on NCPs based on the genomic templates, whereas it was the same when comparing the corresponding free DNA libraries (**Fig. 1e**). Furthermore, *k*-mer enrichment from TF binding to NCPs based on a genomic template was significantly less correlated with that from binding to W601-based libraries than with that from other genomic template libraries (**Fig. 1f,g**). Together, these results revealed that the nucleosomal sequence context influences TF:NCP binding recognition.

### CEBPB exhibits multiple pioneer binding modes to W601-based nucleosomes

We reasoned that the differences in TF pioneer binding to the W601-based PIONEAR-seq libraries (**Fig. 1e**) may be due to the higher stability of W601-based NCPs than of NCPs based on genomic loci^34^. We therefore asked if distinct pioneer binding modes occur on W601-based NCPs as compared to genomic templates. Since the TFs’ DNA binding specificity was the same on NCPs as on free DNA (**Fig. 2a, Extended Data Fig. 3a-c, Extended Data Table 1**), we asked if the TFs may differ instead in their positional preference for binding to W601 versus genomic sequence NCPs.

**Figure 2:**
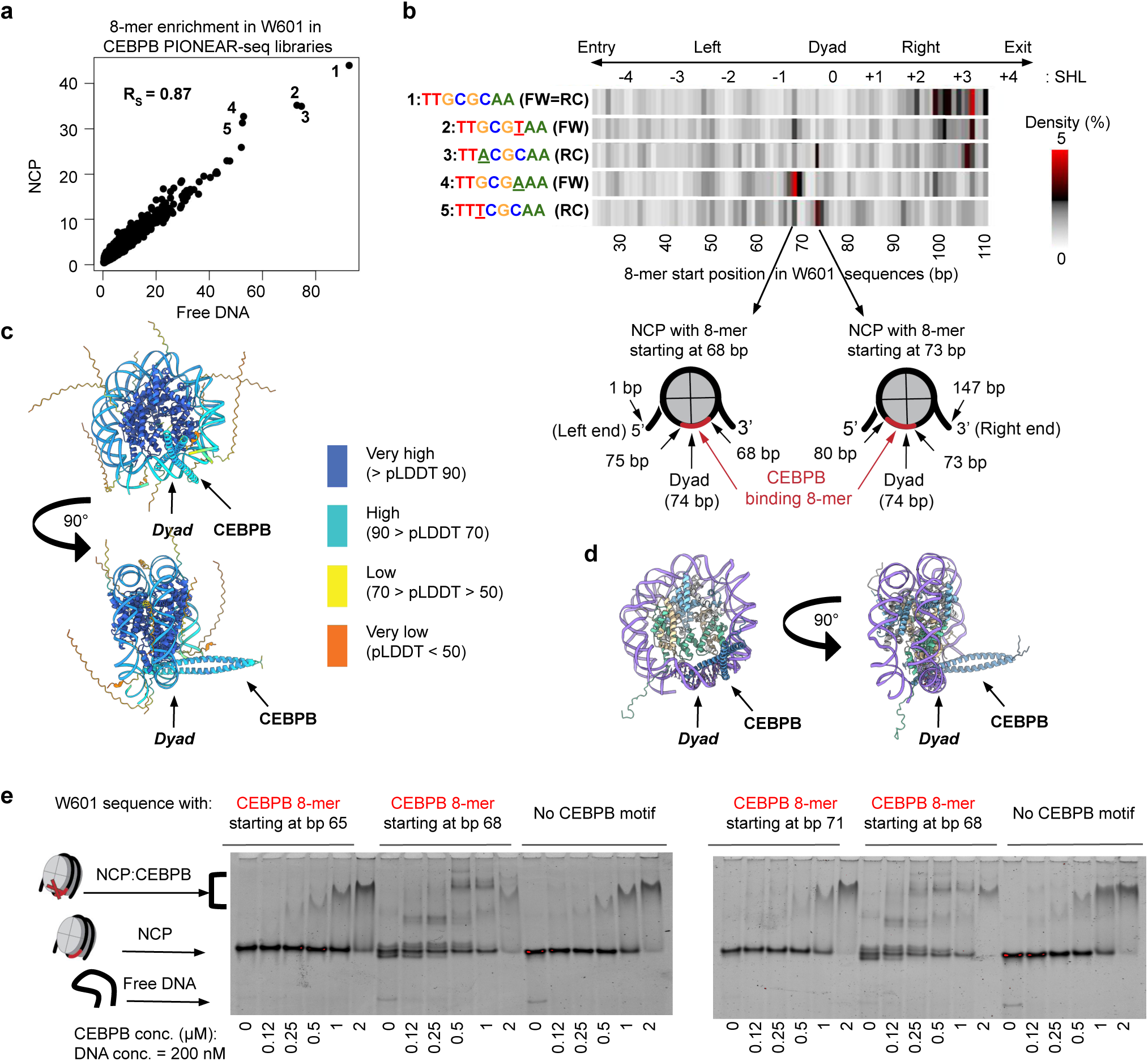
CEBPB recognizes binding sites at specific locations of the W601-based NCPs. **(a)** 8-mer enrichment in W601 libraries bound by CEBPB either as free DNA (x-axis) or NCPs (y-axis) (Spearman R = 0.97). The free DNA library came from Round 1 selection, whose enrichment levels were most similar to those of NCP libraries from Round 4 selection. The most enriched 8-mers are numbered according to the sequences shown in (b). **(b)** Positional density profiles of the indicated 8-mers in the N_20_ tiles of NCP libraries from (a). Bottom: Arrows point to schemas of NCPs containing the CEBPB binding 8-mers at the indicated positions, at position 68-75 bp (left) or 73-80 bp (right) **(c-d)** Structures predicted by AlphaFold3 **(c)** and Rosetta **(d)** of the DNA binding domains of a CEBPB dimer bound to the CEBPB recognition site inserted at position 68-75 bp in a W601 nucleosome. A scale bar showing per-residue confidence metric called predicted local distance difference test (pLDDT) from the AlphaFold3 prediction is shown on the right. **(e)** Representative EMSAs showing CEBPB binding to W601 NCPs carrying a CEBPB binding 8-mer at the indicated positions. The arrows indicate the position of the different configurations of the DNA in the reactions (residual free DNA, reconstituted NCP, CEBPB-bound NCP).

For CEBPB, the most enriched 8-mer (TTGCGCAA) occurred preferentially at specific locations in the right half of W601-based nucleosomes around superhelical location [SHL] +3 (**Fig. 2b**), consistent with a prior study that found that CEBPB binds to the ends of NCPs assembled on fully randomized DNA libraries^14^. Surprisingly, however, we found that the most enriched 8-mers were also enriched at two specific positions near the dyad at approximately SHL −1 to 0 (8-mers spanning positions 68-75 bp and 73-80 bp according to the template coordinate scheme). Furthermore, these two sites differed in their motif orientation: the preferred 8-mers at position 68-75 bp (TTGCG[T/A]AA) were the reverse complement of the 8-mers preferred at 73-80 bp (TT[T/A]CGCAA). Since the expected position of the dyad is at 74 bp, these two peaks represent the same binding site symmetrically positioned on the left and on the right side of the dyad (bottom schemas in **Fig. 2b**). Analysis of binding to the associated free DNA libraries confirmed that these preferential binding locations were specific to nucleosomes (inset, **Extended Data Fig. 3d**).

Previous reports have indicated that CEBPB and, more generally, bZIP and bHLH TFs are unlikely to bind internal nucleosome positions^14^ because of the potential steric clash between their extended scissor-like DNA-contacting alpha-helices and the histone octamer^16,23^. To reconcile these structural concerns with our findings, we created *in silico* models for the CEBPB-nucleosome complex using two orthogonal modeling approaches: a deep learning-based approach using AlphaFold3^47^ and molecular modeling using Rosetta3^48^ (**Supplementary Information**).

Using AlphaFold3, we generated a CEBPB:NCP structure for a W601 NCP containing a CEBPB binding site at position 68-75 bp (**Supplementary Table 6**). This structure predicts that the C-terminal bZIP domain of CEBPB (CEBPB CTD) in a dimer clamps onto the CEBPB binding site near the dyad, with one CEBPB CTD threading through the space between the dyad and the globular domain of histone H3 (**Fig. 2c**). The overall mode of binding resembles the CEBPB CTD dimer recognizing a linear CEBPB consensus sequence^49^, although the CEBPB binding site adopts a slight curvature in the nucleosomal environment. In this binding mode, the position and spacing of the CEBPB binding site near the dyad are crucial for CEBPB recognition. Even a slight shift of DNA can significantly impact the CEBPB binding mode by occluding the small space between the dyad and the histone H3 globular domain, preventing one CEBPB CTD from threading through. When a 3-bp shift of the CEBPB motif from the 68-75 bp position was introduced, AlphaFold3 exhibited: 1) lower predicted alignment error (PAE) scores between the CEBPB dimer and the histones or nucleosomal DNA, and 2) the absence of perpendicularly docked topology of CEBPB seen in the unshifted complex structure (**Extended Data Fig. 4**). These observations indicate that the insertion of the CEBPB motif at the 68-75 bp position enables CEBPB to recognize the nucleosome in a unique conformation, and even a 3-bp shift of this motif may disrupt this binding mode. Altogether, these modeling results support the preferential position CEBPB may need to recognize its cognate sequence near the dyad, as revealed by our PIONEAR-seq results (**Fig. 2b**).

Using Rosetta3, we employed a comparative modeling strategy as an orthogonal approach to obtain a CEBPB:NCP structure. Briefly, we used a previously solved nucleosome crystal structure (PDB: 1AOI) as a template and replaced the nucleosomal DNA sequence with the 146 bp W601 DNA sequence containing a CEBPB-binding 8-mer at position 68-75 bp predicted by PIONEAR-seq (**Fig. 2b**). Then, we superimposed the CEBPB CTD dimer-DNA complex (PDB: 6MG1)^49^ on the nucleosome, combined the pre-positioned CEBPB:DNA and nucleosome structures, and relaxed the whole complex using Rosetta3’s Relax application^50^. Overall, the CEBPB structure from Rosetta3 was in good agreement with the AlphaFold3 structure (**Fig. 2d**).

To corroborate these findings biochemically, we used electrophoretic mobility shift assays (EMSAs) to compare binding by CEBPB to W601 nucleosomes carrying a CEBPB binding sequence at different positions (**Supplementary Table 7**). The results showed that CEBPB bound specifically to W601-based nucleosomes with a CEBPB binding sequence at position 68-75 bp, whereas CEBPB association with NCPs with the binding site at position 65-72 bp or 71-78 bp was indistinguishable from that with NCPs lacking a CEBPB binding sequence (**Fig. 2e, Extended Data Fig. 5**). These results support our PIONEAR-Seq results and support our structural model prediction that a slight shift of DNA can significantly impact the CEBPB binding mode.

Altogether, these results revealed that CEBPB, which was reported previously to be an “end binder”^14^, can also bind at specific positions near the dyad in W601-based NCPs.

### Sequence context regulates positional preferences in pioneer binding

To investigate the potential mechanisms underlying the multiple binding modes of CEBPB in the W601 library (**Fig. 2**), we first asked whether they were a property specific to CEBPB or alternatively to the W601 NCPs. Inspection of the positional preferences of other TFs that exhibited pioneer binding by PIONEAR-seq (*i.e.*, OCT4, RUNX2, PAX7, FOXA1, GATA1, GATA4) (**Fig. 1b**) revealed that similar to CEBPB, RUNX2 and PAX7 also showed strong preferences for binding their recognition sequences at specific internal positions of the W601 NCPs (**Fig. 3a**), in contrast to their lack of positional binding preferences, as expected, in the corresponding free DNA libraries (**Extended Data Fig. 3d**). PAX7’s most enriched 8-mers (AAT[T/C]GATT; **Extended Data Fig. 3b**) were found mostly at specific positions proximal to the dyad, while those of RUNX2 (TTGCGGTT; **Extended Data Fig. 3c**) were bound preferentially at 10-bp periodicity in the right half of W601-based NCPs (**Fig. 3a**), consistent with a prior study that found RUNX3 to bind with the same periodicity on NCPs assembled on fully degenerate sequence^14^.

**Figure 3:**
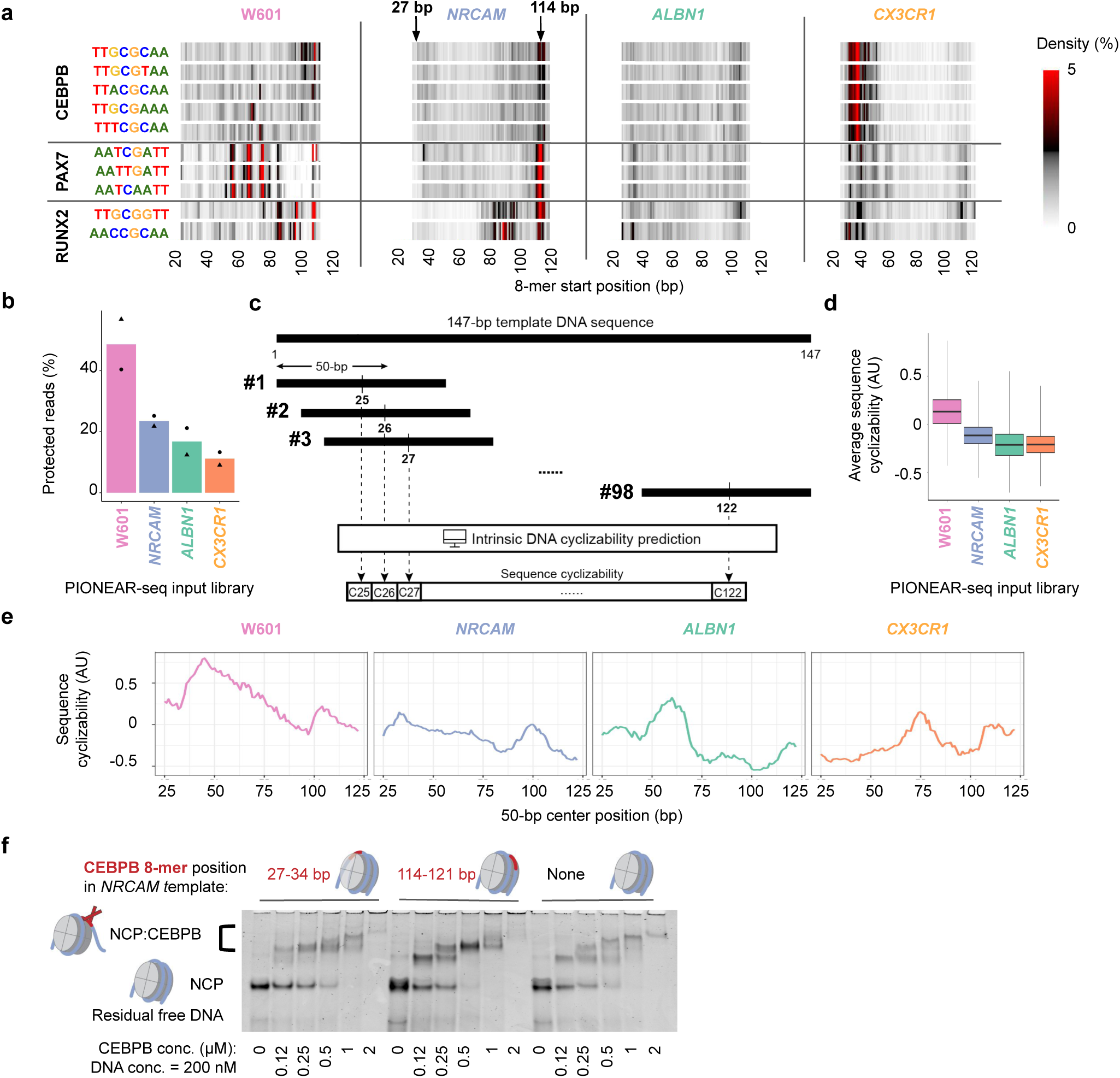
The DNA composition of nucleosomes modulates the positional preferences of TF binding. **(a)** Positional density profiles of the most enriched 8-mers for CEBPB, PAX7, RUNX2 in the N_20_ tiles of NCP libraries based on different templates. Arrows on the top of *NRCAM* profiles highlight an example of 8-mer positions that are symmetrical around the dyad, but show different levels of motif enrichment, suggesting that the TFs prefer to bind at 114 bp more than at 27 bp. **(b)** Percentages of sequences from each of the W601, *NRCAM, ALBN1, CX3CR1* NCP libraries after MNase digestion and sequencing (2 replicates). Negative controls performed without MNase resulted in sequences from each of the 4 libraries at ∼25% of the total pool (**Extended Data Fig. 8a**). **(c)** Workflow for the prediction of DNA cyclizability along 147-bp sequences using a deep learning model trained on 50-bp loop-seq data. DNA sequence is converted to 50-bp sliding windows and mapped to a cyclizability score vector with a prediction model. In the case of a 147-bp sequence, the 50-bp sliding windows creates a vector of 98 cyclizability scores. **(d)** For the indicated template, the distributions of the average sequence cyclizability in the associated input libraries. **(e)** Sequence cyclizability profiles of the indicated templates for each position of the 50-bp sliding windows. **(f)** Representative EMSA showing CEBPB binding to *NRCAM* NCPs carrying the CEBPB motifs at the indicated positions. The arrows indicate the positions of the different configurations of the DNA in the reactions (residual free DNA, reconstituted NCP, CEBPB-bound NCP).

In contrast, the dyad and periodic binding modes exhibited by these TFs on W601-based NCPs were absent from the NCPs based on genomic sequences (**Fig. 3a**). Moreover, we noticed that for these 3 TFs the binding occurred preferentially at the 3’ end of *NRCAM*-based nucleosomes and at the 5’ end of CX3CR1-based nucleosomes, while in *ALBN1* libraries there was no apparent positional preference (**Fig. 3a**), suggesting that the positional preference was not an inherent property of their DNA binding domains, but rather varied depending on the sequence context of the nucleosomal DNA.

We reasoned that W601-specific preferences for dyad and periodic binding by CEBPB, PAX7, and RUNX2 might originate from the higher stability of W601-based nucleosomes^18^ as compared to nucleosomes based on genomic sequences, which might permit greater end breathing due to a looser structure, as suggested by a direct comparison between W601 and ALBN1 nucleosome structures^27^. To investigate such potential sequence-dependent differences in DNA accessibility across the different nucleosomes in our survey, we assayed the input NCP libraries by MNase-seq (**Supplementary Information**). The MNase-seq profiles showed that the W601 NCPs were the most protected from MNase digestion and therefore provided the least accessible ends for TF binding (**Fig. 3b, Extended Data Fig. 6a**).

### DNA flexibility underlies nucleosomal end-binding preferences of TFs

A prior single-molecule study found that the sequence composition of the inner quarters of W601-based nucleosomal DNA affects the dynamics of the adjacent outer quarters^30^. These authors propose a model in which the inherent DNA rigidity due to its sequence is less favorable for wrapping around the octamer, thus exposing potential TF binding sites. We therefore asked whether DNA flexibility might control the sequence-dependent end-binding preferences that we observed in our PIONEAR-seq data. To analyze DNA flexibility of nucleosomes, we used two deep learning models^51,52^ trained on loop-seq data^53,54^, which quantifies the ability of ∼100-bp DNA sequences to form loops *in vitro* (’*cyclizability*’) as a proxy for the DNA flexibility. We used these deep learning models, which predict the sequence cyclizability of user-input 50-bp sequences, to score 50-bp sliding windows tiling across the nucleosomal sequences (**Fig. 3c**). Comparison of the average cyclizability of the DNA input libraries based on the different templates revealed that the W601 library was overall much more flexible than libraries based on the genomic sequences (**Fig. 3d**). Intriguingly, the NCP input libraries showed the same trend for DNA cyclizability as for protection from MNase digestion (**Fig. 3b,d**), suggesting that the flexibility of the nucleosomal DNA sequence controls the accessibility of the associated nucleosomes.

We next examined the cyclizability profiles along the templates. We found that the predicted cyclizability was nonhomogenous across the templates, with a tendency to decrease from the 5’ to the 3’ end for the W601 and *NRCAM* templates and from the 3’ to the 5’ end for the *CX3CR1* template (**Fig. 3e**). Highly similar cyclizability profiles were obtained using both deep learning models (**Extended Data Fig. 6b**). Strikingly, we noticed that the preferred positions for TF pioneer binding tended to occur at the less cyclizable side (compare **Fig. 3a** and **Fig. 3e**). Examination of the cyclizability of the DNA input library confirmed that the inserted N_20_ random tiles did not alter the asymmetry trend of sequence cyclizability (**Extended Data Fig. 6c**). A prior observation that the end of W601-based nucleosomes adjacent to more rigid internal DNA unwraps more easily^30^ led us to hypothesize that such asymmetry in DNA flexibility might drive TF binding preferentially to the end of nucleosomes adjacent to the more rigid internal DNA. To test this hypothesis, we used EMSAs to compare CEBPB binding to *NRCAM* NCPs built with CEBPB motifs positioned on the more flexible end (at position 27-34 bp) versus the more rigid end (at position 114-121 bp) (**Supplementary Table 7)**. In agreement with our predictions, we observed stronger CEBPB binding to NCPs carrying the motif on the more rigid end of the *NRCAM* nucleosomal DNA (**Fig. 3f, Extended Data Fig. 6d-f**).

These flexibility-associated end-binding preferences were either weaker or absent for the other four TFs (OCT4, FOXA1, GATA1, GATA4) that bound nucleosomes by PIONEAR-seq. In the case of OCT4, which showed marked end binding in libraries based on both W601 and genomic templates, the bound end was defined by the motif orientation (the forward motif AATT[A/T]GCTAT at the 5’ end and the reverse complementary motif TATGC[A/T]AATT at the 3’ end; **Extended Data Fig. 7a**), in agreement with prior *in vitro* studies of OCT4 binding to nucleosomes based on W601^10^ and genomic sequences^4^. FOXA1 also displayed end binding preferences across all the NCP libraries (**Extended Data Fig. 7b**), in agreement with a prior study that found FOXA2 to be an end binder^14^. Finally, the binding modes of GATA1 and GATA4, which showed very similar binding site positional preferences along the nucleosomes (**Extended Data Fig. 7c,d**), were highly consistent with those of CEBPB, PAX7 and RUNX2 on W601, *NRCAM* and *CX3CR1* templates, and were consistent with a prior study that found GATA3 dimers to bind tandem GAT sites on W601-based nucleosomes^12^; however, the associated free DNA libraries exhibited enrichment of GATA binding sites at several positions, which biased the interpretation of their NCP binding profiles. Overall, the positional preference results from the prior Zhu *et al.* survey of TF binding to nucleosomes containing 101-bp randomized sequence^14^ were in general agreement, with some exceptions, with our PIONEAR-Seq results for TF binding to sites on W601-based nucleosomes; larger differences, however, were apparent when comparing the binding modes called by Zhu *et al.* with the binding modes that we observed for TF interactions with NCPs assembled on genomic nucleosome positioning sequences (**Extended Data Fig. 8; Supplementary Discussion**).

Altogether, these results revealed that certain TFs use different binding modes to pioneer nucleosomes and that those modes are regulated by the sequence context in which their cognate motif is embedded. We next asked if DNA flexibility controls nucleosome binding by TFs beyond those that we assayed by PIONEAR-Seq. Here, we analyzed published NCAP-SELEX data for 135 human TFs for binding to NCPs that included 101-bp internal, randomized sequence (N_101_) (**Supplementary Information**; **Fig. 4a**, **Extended Data Fig. 9a; Supplementary Table 8**). For each TF, we inspected NCPs containing TF binding sites at the ends of the randomized regions (’end binding reads’) (**Fig. 4a, Extended Data Fig. 9a)**. For a simplified representation of the end binding reads, we re-oriented the sequencing reads such that the TF binding sites are always on the left (’re-orientation’, **Fig. 4a**) and then profiled their cyclizability as described above (**Fig. 3c**).

**Figure 4:**
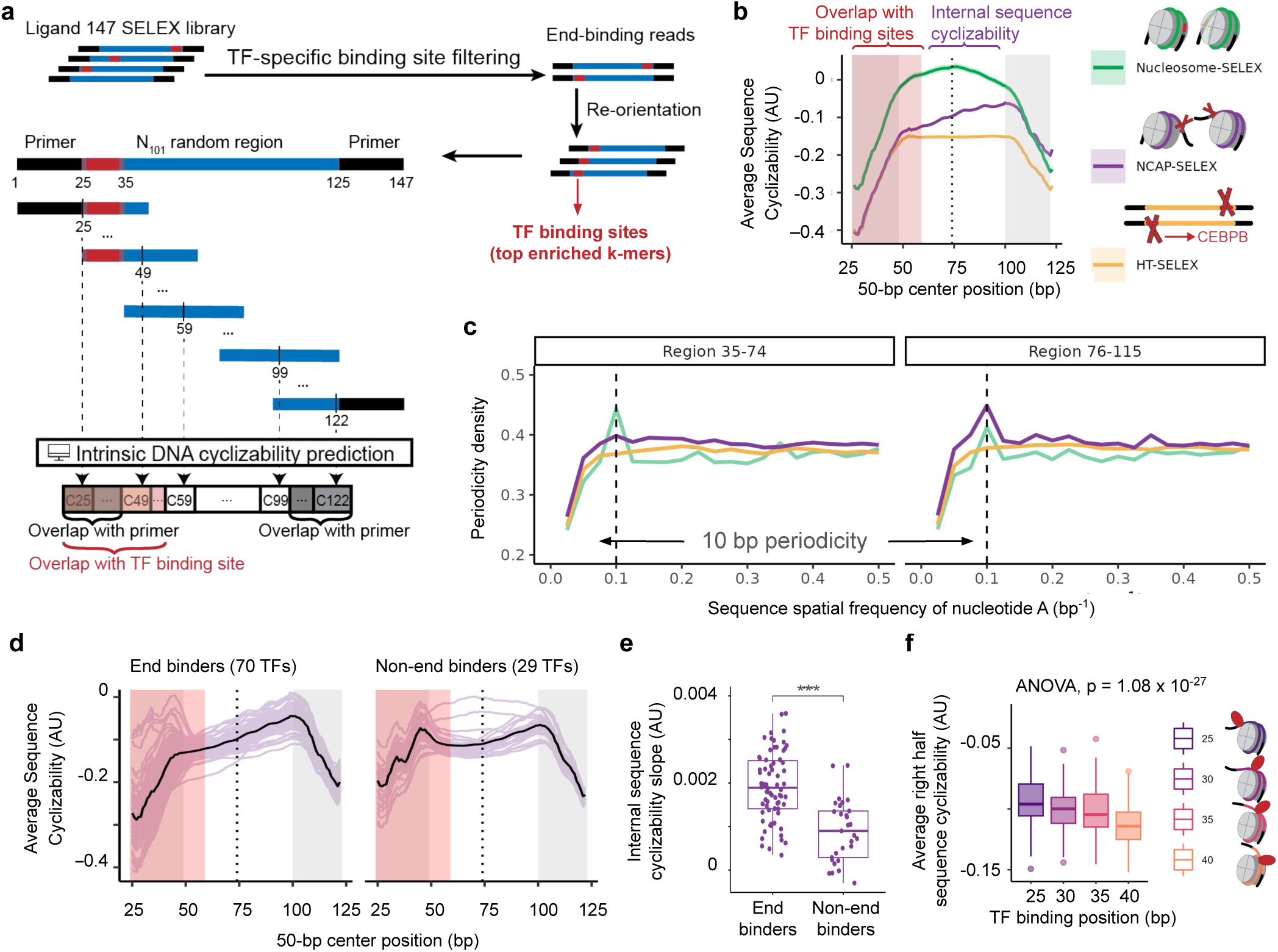
DNA sequence composition drives TF binding towards the less flexible end of the nucleosomes. **(a)** Workflow of the cyclizability analysis of NCAP-SELEX, HT-SELEX, and Nucleosome-SELEX data. Each library consists of 147-bp reads with a 101-bp random region (N_101_, blue), flanked by fixed primers at the end (black). Reads containing the most (*i.e.*, top 10) enriched *k*-mers (red) in the first or last 10 base pairs of the random region are filtered, re-oriented to have the binding site on the left side, and then profiled for sequence cyclizability using the 50-bp sliding window model described in Fig. 3c. **(b)** Averaged profiles of sequence cyclizability in Nucleosome-SELEX (green), CEBPB NCAP-SELEX (purple) and CEBPB HT-SELEX libraries (yellow). The 10 most enriched *k*-mers (and their reverse complements) found in CEBPB NCAP-SELEX were used to filter the ‘end-binding’ reads in the 3 libraries. **(c)** Averaged 10-bp periodicity intensity (dashed line) of nucleotide ‘A’ across filtered reads from the Nucleosome-SELEX library (green), the CEBPB NCAP-SELEX library (purple), the CEBPB HT-SELEX library (yellow), respectively. The internal random regions without the TF binding sites are separated into two symmetrical regions on the sequence, adjacent to the TF binding sites (left, region 35-74 bp) and distant from the TF binding sites (right, region 76-115). **(d)** Bulks of averaged profiles of sequence cyclizability for TFs classified as end binders (left) or non-end binders (right) from a published NCAP-SELEX screen^14^. For each TF, the averaged profile of sequence cyclizability was obtained as depicted in (a,b). **(e)** Fitted linear regression slopes on the internal sequence cyclizability of NCAP-SELEX data for end binding versus non-end binding TFs (Wilcoxon Rank Sum Test, *P* = 2.3 x 10^-7^). **(f)** Averaged cyclizability of the right half sequences of NCAP-SELEX libraries for 77 end-binding TFs. Data were stratified according to the distance between the TF binding sites and the 5’ end of the nucleosome (**Extended Data Fig. 9f**) and the distance was used to test a linear regression model by ANOVA (*P* = 1.08 x 10^-27^).

We found that CEBPB end binding sites in the NCAP-SELEX data tended to occur on the more rigid side of the nucleosomes (purple curve, **Fig. 4b**), consistent with our PIONEAR-Seq data on CEBPB end-binding preferences on *CX3CR1, NRCAM* and *W601* nucleosomes (**Fig. 3a,e,f)**. As negative controls, analysis of neither free DNA selected for binding to CEBPB alone (“HT-SELEX”) nor DNA selected for binding to nucleosomes alone (“nucleosome-SELEX”, **Extended Data Fig. 9a**) showed a difference in the DNA cyclizability of the nucleosome ends with versus without CEBPB binding sites (**Fig. 4b**). Notably, when compared to CEBPB HT-SELEX reads, the cyclizability of N_101_ in nucleosome-SELEX reads was much higher, indicating that they are overall more flexible, as expected for sequences selected to form nucleosomes (**Fig. 4b**).

Strong nucleosome positioning sequences have been observed to display ∼10-bp nucleotide periodicity^55^. We found that the nucleotide periodicity in the left (35-74 bp) and right (76-115 bp) sides of the N_101_ regions of the nucleosomes were highly concordant with the cyclizability analysis: the N_101_ sides adjacent to the binding sites (left sides) did not show any 10-bp periodicity, although the N_101_ sides located distal to the CEBPB binding sites (right sides) exhibited pronounced 10-bp periodicity (purple curves, **Fig. 4c**), suggesting that the CEBPB-bound sides are less likely to be associated strongly with histone octamer. In contrast, the TF-unbound nucleosome-SELEX controls exhibited strong 10-bp periodicity in both halves, in agreement with their expected preference to form nucleosomes (green curves, **Fig. 4c**), whereas the HT-SELEX free DNA controls did not show 10-bp periodicity, as expected (yellow curves, **Fig. 4c**).

To aid in generalizing these results to TFs beyond CEBPB, we assigned two scores to quantify each TF’s cyclizability profile: (1) the average cyclizability of the internal NCP region as a proxy for its overall flexibility, and (2) the slope of the internal region as a proxy for its differential flexibility between the two ends (**Extended Data Fig. 9b).** We then classified the 137 TFs assayed by both NCAP-SELEX and HT-SELEX with sufficient read depth for this cyclizability analysis (**Supplementary Information)** into 70 end binders and 29 non-end binders (and 38 undetermined) (**Extended Data Fig. 9c**). The results revealed that the 70 end binders were enriched for binding their sites on the more rigid ends of the nucleosomes as compared to the 29 non-end binders (*P* = 2.3 x 10^-7^, Wilcoxon rank sum’s test) (**Fig. 4d,e**). In contrast, the HT-SELEX (and TF-unbound nucleosome-SELEX) controls did not show a significant difference between the end versus non-end binders (**Extended Data Fig. 9d**). Moreover, we found that the end binders exhibited the same 10-bp dinucleotide periodicity pattern as did CEBPB (compare **Fig. 4c** and **Extended Data Fig. 9e**), further supporting our model that the TF end-binding mode is regulated by the broader sequence context of the nucleosome and favored on sites located at the more rigid side of nucleosomes.

To investigate whether the DNA flexibility of the nucleosome affects the recruitment of TFs at more internal sites, we stratified the TF-bound NCP reads according to the position of the TF binding sequence within the N_101_ regions into 4 subsets: 25-35, 30-40, 35-45, 40-50 bp from the left end of the reads (**Supplementary Information)**. We found that the end binding of CEBPB extended to more internal positions if they were flanked by more rigid DNA regions towards the dyad (left, **Extended Data Fig. 9f**). This trend generalized to all 70 end-binders, with the average cyclizability of the right side sequences decreasing proportionally to the extent of the internal binding by the TF (*P* = 1.08 x 10^-27^, ANOVA) (**Fig. 4f**, **Extended Data Fig. 9f**).

To determine if DNA flexibility modulates TF binding to nucleosomes *in vivo*, we analyzed TF genomic occupancy (ChIP-seq) and nucleosome occupancy (MNase-seq) data from human K562 cells^56^. From those data, we identified 11,567 nucleosomes that overlapped a CEBPB ChIP-seq peak and contained a CEBPB binding site within 30 bp of one end of the nucleosomes (**Supplementary Information**, **Fig. 5a, Supplementary Table 9**). We found that these 11,567 CEBPB-bound nucleosomes were significantly more rigid (*i.e*., had lower cyclizability) in the internal region flanking the CEBPB motif than nucleosomes that contained the CEBPB binding site but were not bound by CEBPB (**Fig. 5b**).

**Figure 5:**
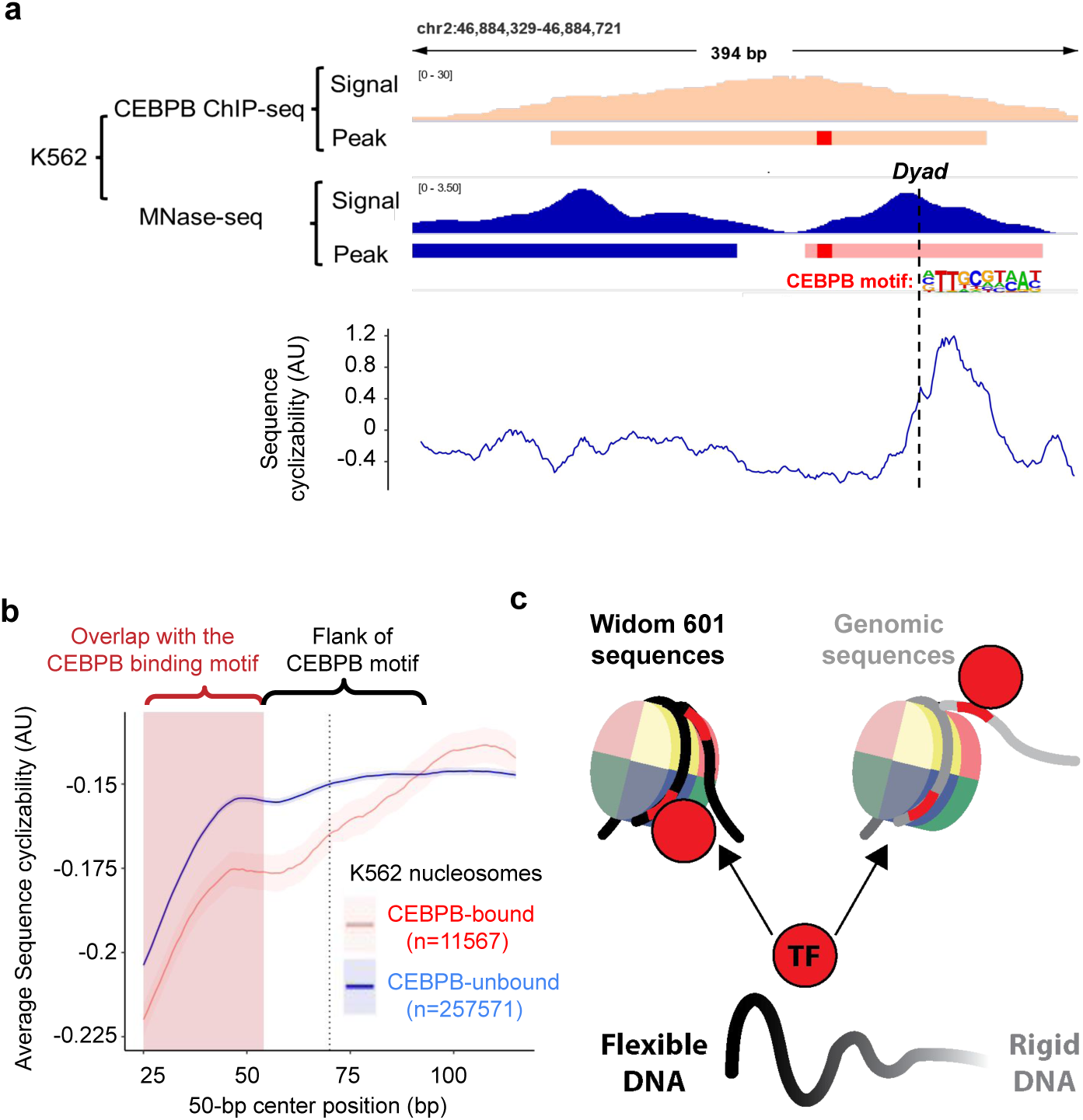
Sequences of *in vivo* CEBPB-bound nucleosomes shows the DNA flexibility asymmetry predicted by *in vitro* CEBPB-bound nucleosomes. **(a)** CEBPB ChIP-seq signals and peaks, MNase-seq signal and nucleosome calls in K562 cells (top tracks) and genome sequence cyclizability computed by a 50-bp sliding window model as in Fig. 3c (bottom track) across the indicated genomic region (chr2: 46,884,329-48,884,721). **(b)** Nucleosomes in K562 cells that contained CEBPB motifs within 30-bp from the ends were divided into two groups according to their overlap with CEBPB ChIP-seq peaks. Average profiles of sequence cyclizability for the two groups were computed as in Fig. 4b. The cyclizability of the 50-bp window centered at 55 bp from the nucleosome end with the CEBPB binding motif is significantly different between CEBPB-bound nucleosomes and nucleosomes not bound by CEBPB (“CEBPB-unbound”). **(c)** Graphical representation of the role of DNA flexibility in regulating positional preferences for TF binding to nucleosomes.

## Discussion

In this study, we developed PIONEAR-seq, a high-throughput biochemical assay for analysis of the effects of nucleosomal DNA composition on pioneer binding, which we used to analyze the DNA sequence specificity and positional preference of binding by seven pioneers to nucleosomes assembled on four different sequence contexts. In contrast to prior models that have suggested that the positional preference for engaging with nucleosomes (*e.g.*, dyad, periodic, end binding) is an intrinsic property of a particular TF or TF family^8,14,16^, we found that the positional preferences of pioneer binding can also be regulated by the DNA composition of the whole nucleosome. Pioneers bound preferentially to internal sites on high-stability nucleosomes through dyad and periodic binding. The W601 sequence is known to form more stable nucleosomes than do genomic sequences^27,34^, which could minimize ‘end breathing’ of W601 NCPs. Indeed, in our assay W601-based NCPs favored TF binding at more internal regions, such as the ‘single-gyre’ dyad or interspersed sites that followed the 10-bp periodicity of DNA wrapped around the histone octamer^9^. In contrast, TFs preferentially bound the ends of genomic nucleosomes with less flexible DNA. Our results support a model in which the more rigid half of the nucleosomal DNA is more likely to dynamically unwrap, exposing binding sites for TF recognition (**Fig. 5c**).

Our results for TF binding to nucleosomes based on genomic sequences may more accurately reflect the *in vivo* binding preferences of pioneers. Nucleosomes have been found to unwrap their ends in a dynamic way that depends on the local flexibility of the DNA sequence, with unwrapping at one end stabilizing the other end of the nucleosome^30^. Thus, the DNA sequence of a nucleosome can amplify differences in local DNA flexibility into asymmetric nucleosome unwrapping, thus providing a mechanism for regulating TF access to specific loci that is consistent with what we found for TF binding to nucleosomes based on the *CX3CR1* and *NRCAM g*enomic sequences.

The idea of pioneers as a subgroup of TFs with the distinct ability to recognize their cognate DNA motif on nucleosomes remains controversial. A recent study challenged this model by showing that both the archetypal pioneer FOXA1 and its associated non-pioneer HNF4 can ‘pioneer’ for each other *in vivo*, but with different strengths and cooperatively^57^. Furthermore, prior *in vivo* studies have shown that even the most well-established pioneers bind only subsets of genomic occurrences of their recognition sites^32,58,59^. Altogether, these results suggest that instead of a categorical distinction between pioneers and non-pioneers, TF binding to nucleosomes forms a continuum and depends on additional cues. By showing that the DNA sequence of nucleosomes restricts which motif instances are eligible for pioneer binding, we present one potential mechanism for the observed selectivity of motif utilization *in vivo*.

Our results suggest that local DNA flexibility represents another layer of *cis*-regulatory encoding in the genome, going beyond TF binding site motifs. Improved tools that include scoring of DNA sequence flexibility may lead to more accurate prediction of functional TF genomic binding sites and thus aid in the interpretation of noncoding variants. Future studies are needed to determine the influence of epigenomic features and interactions with cofactors and chromatin remodelers on TF binding to nucleosomes and gene regulation.

## Acknowledgements

This work was supported in part by an American Heart Association postdoctoral fellowship (#826614 to K.L.) and by grants from the U.S. National Institutes of Health (R01 HG012246 and R21 HG009268 to M.L.B, R37 GM62437 to P.C.). We thank Taekjip Ha, Park Jonghan, and Sophia Yan for sharing their unpublished software to predict DNA sequence flexibility. We thank members of the Bulyk lab for helpful discussion, M. Wu, R. Kingston and J. Cochrane for technical assistance in the production of octamers and nucleosomes in the initial stages of this project.

## Author Contributions

L.M. and M.L.B. designed the experiments. L.M. performed experiments. L.M. and X.L. performed analyses. L.M., K.L., X.L., and S.S.G. performed molecular modeling and structure interpretation. K.L. and P.A.C. contributed to production of histone octamers and purification of CEBPB. All authors interpreted the results. L.M., X.L. and M.L.B. wrote the manuscript. All authors read and edited the manuscript.

