## Extended Data Figures for "DNA flexibility regulates transcription factor binding to nucleosomes"

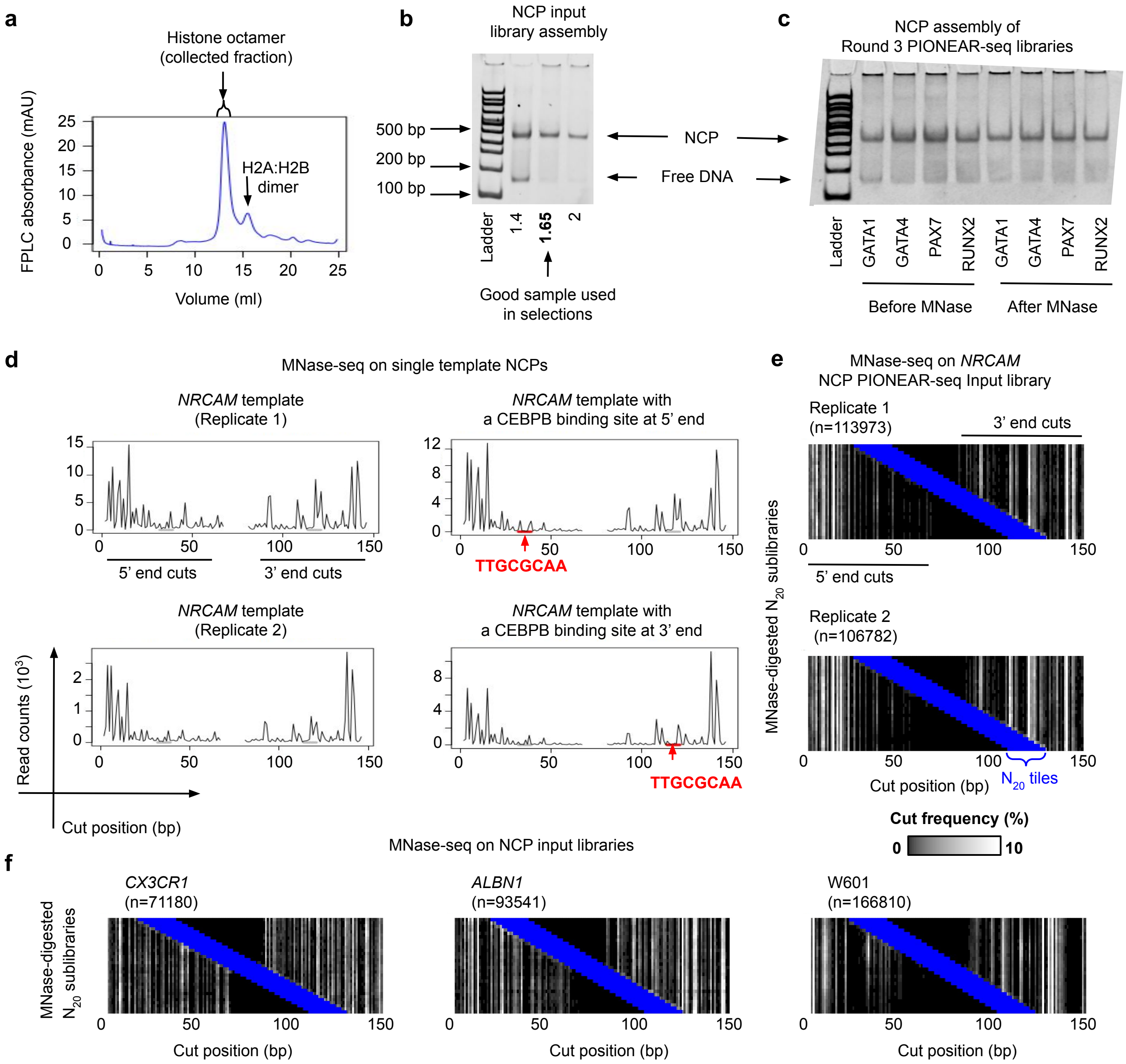


**Extended Data Fig. 1**

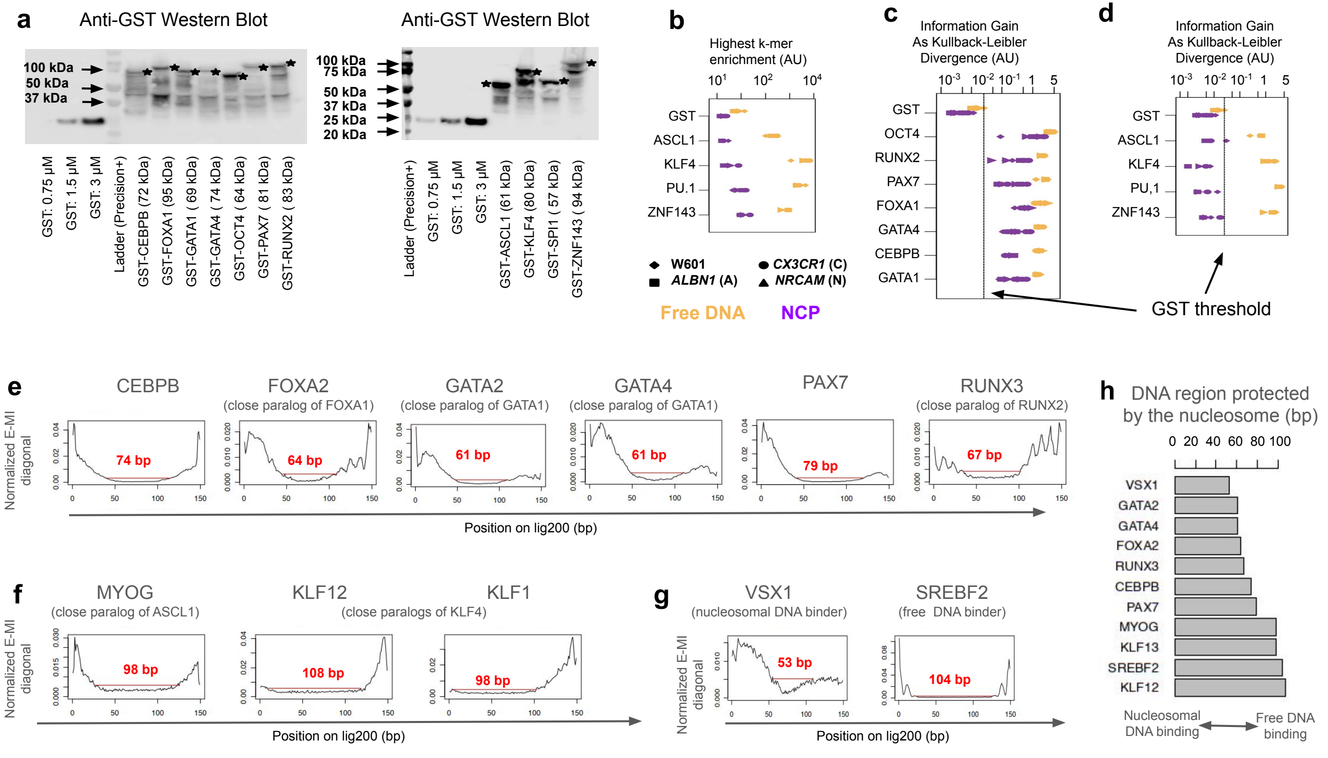


**Extended Data Fig. 2**

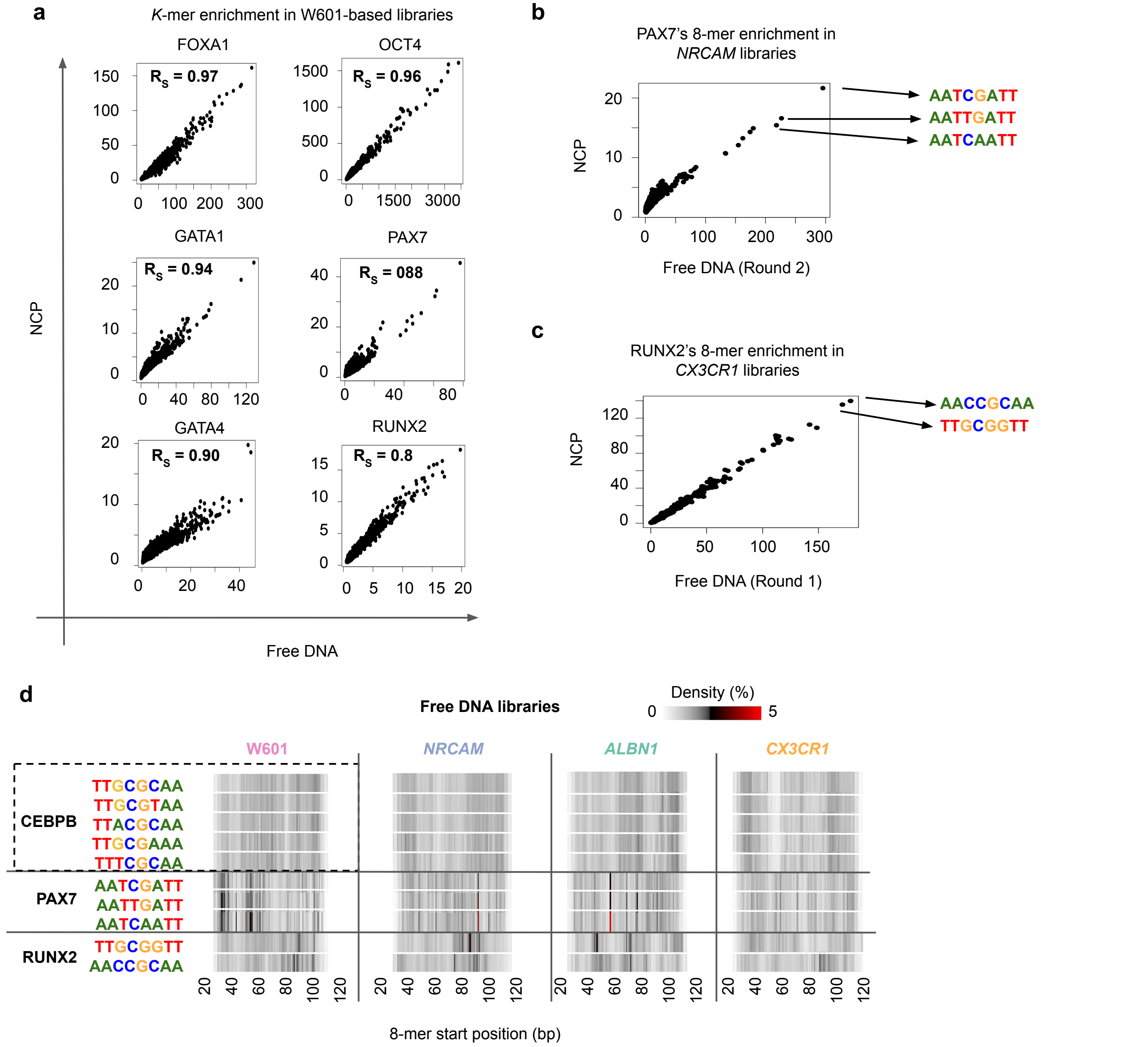


**Extended Data Fig. 3**

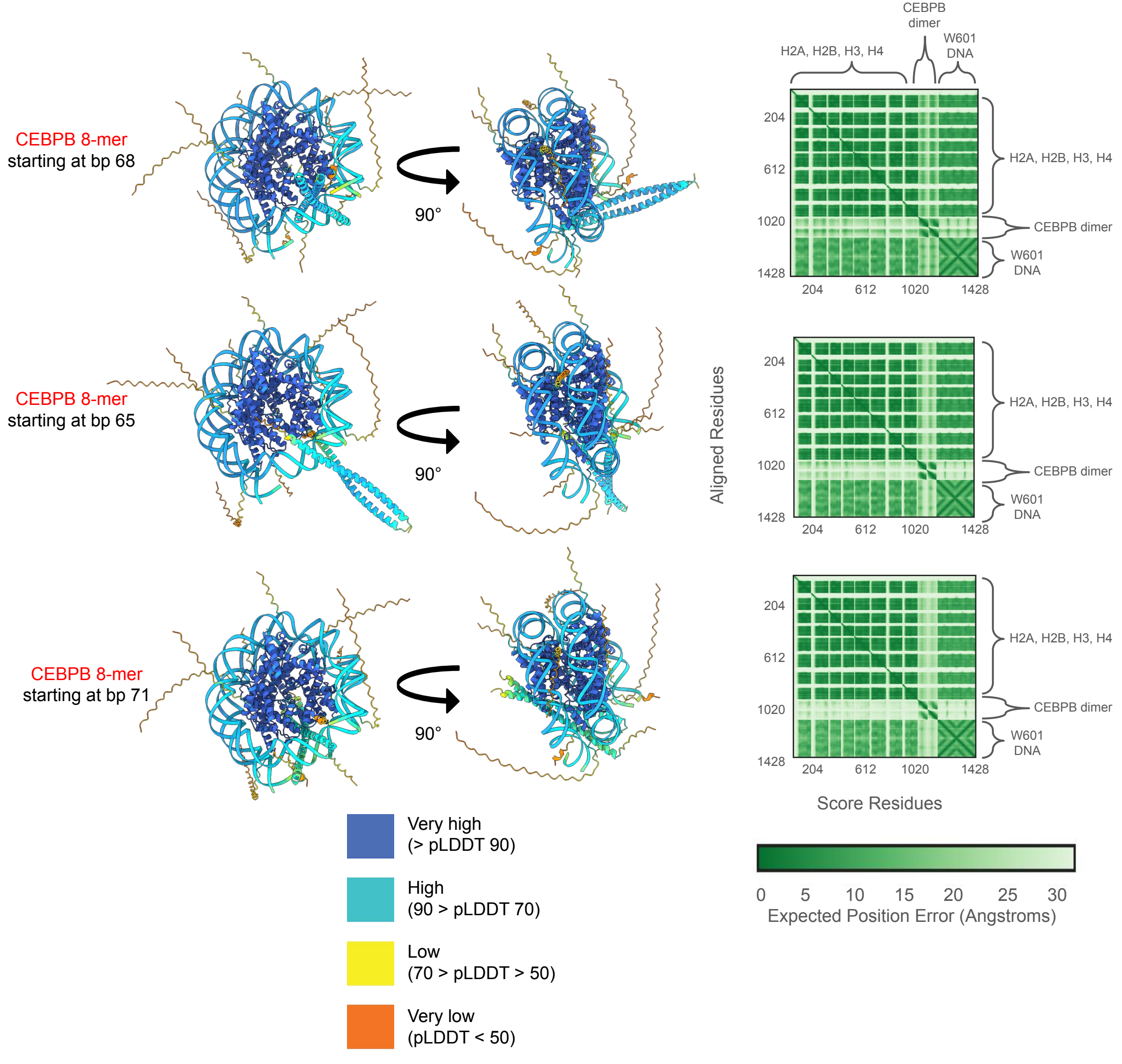


**Extended Data Fig. 4**

### Replicate of EMSAs in Fig. 2e

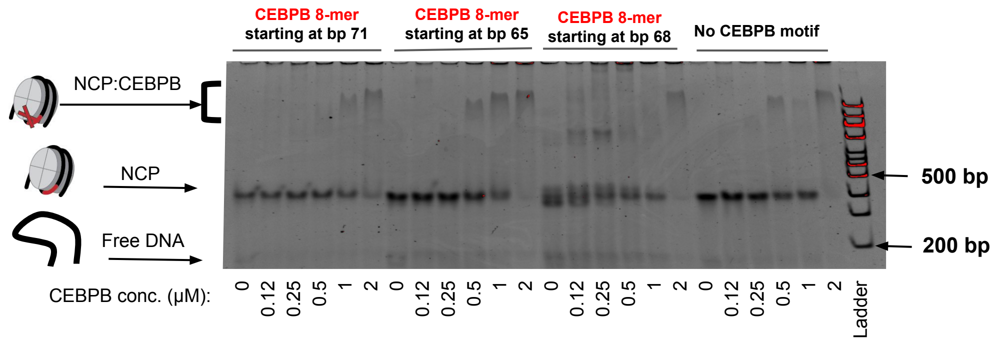


### Same EMSAs as in Fig. 2e, but with free DNA instead of NCPs

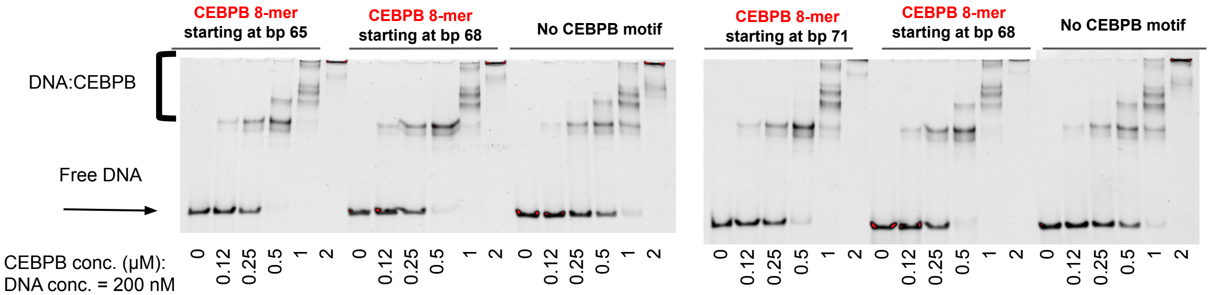


### EMSA for full-length CEBPB binding to W601-based NCPs

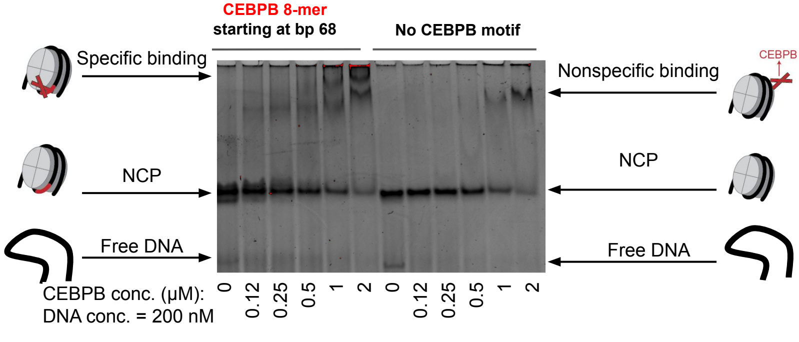


**Extended Data Fig. 5**

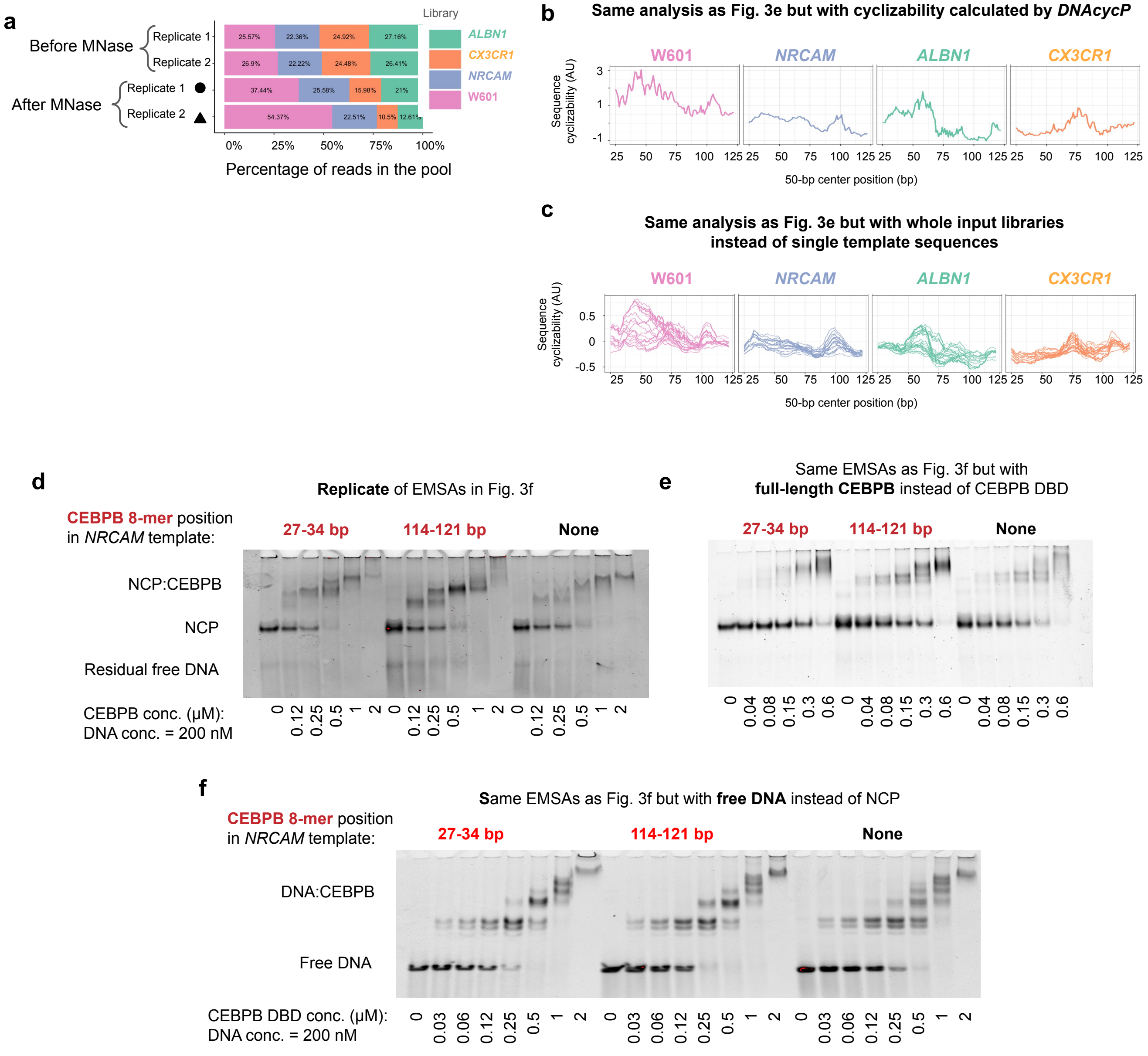


**Extended Data Fig. 6**

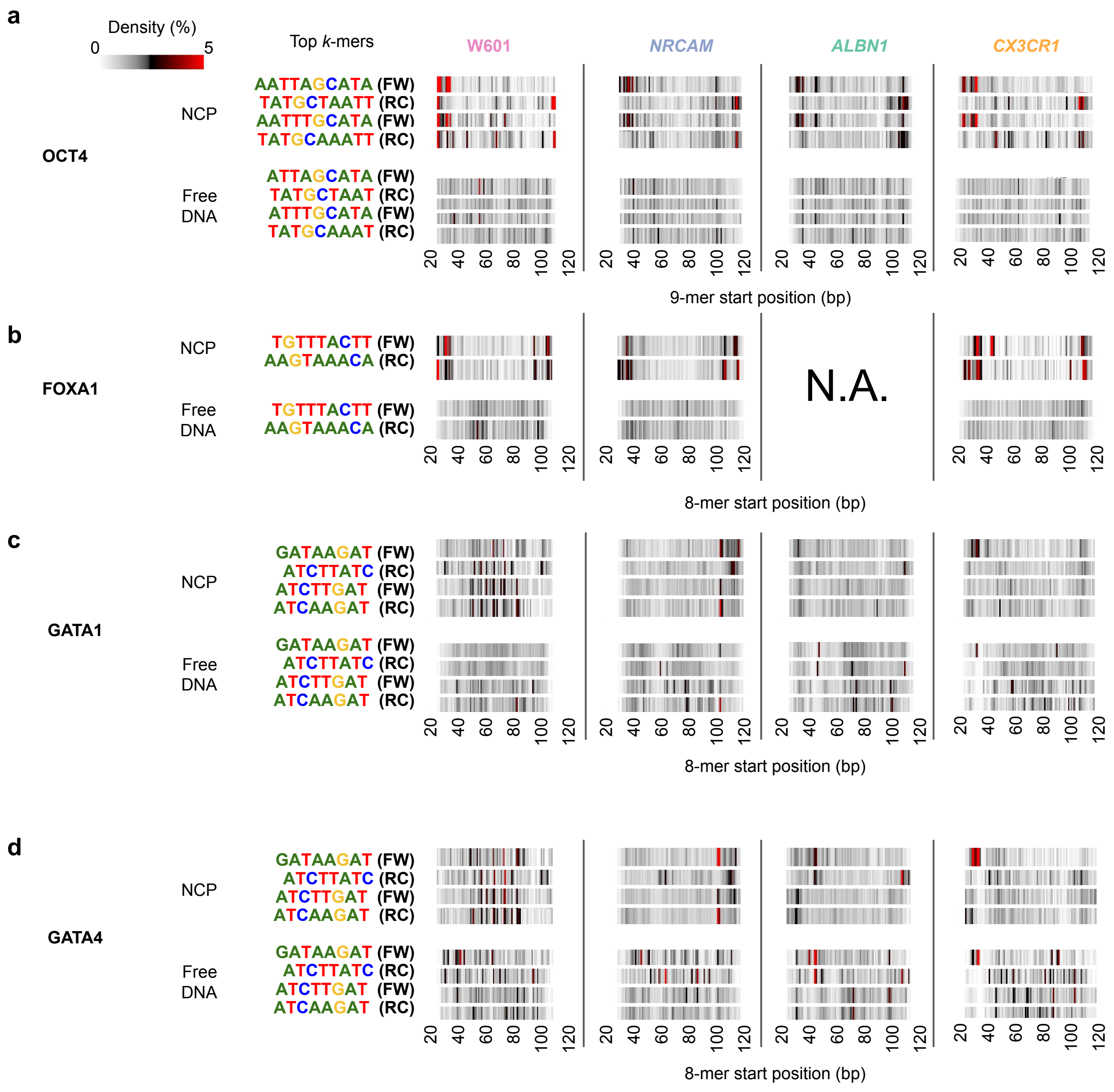


Extended Data Fig. 7

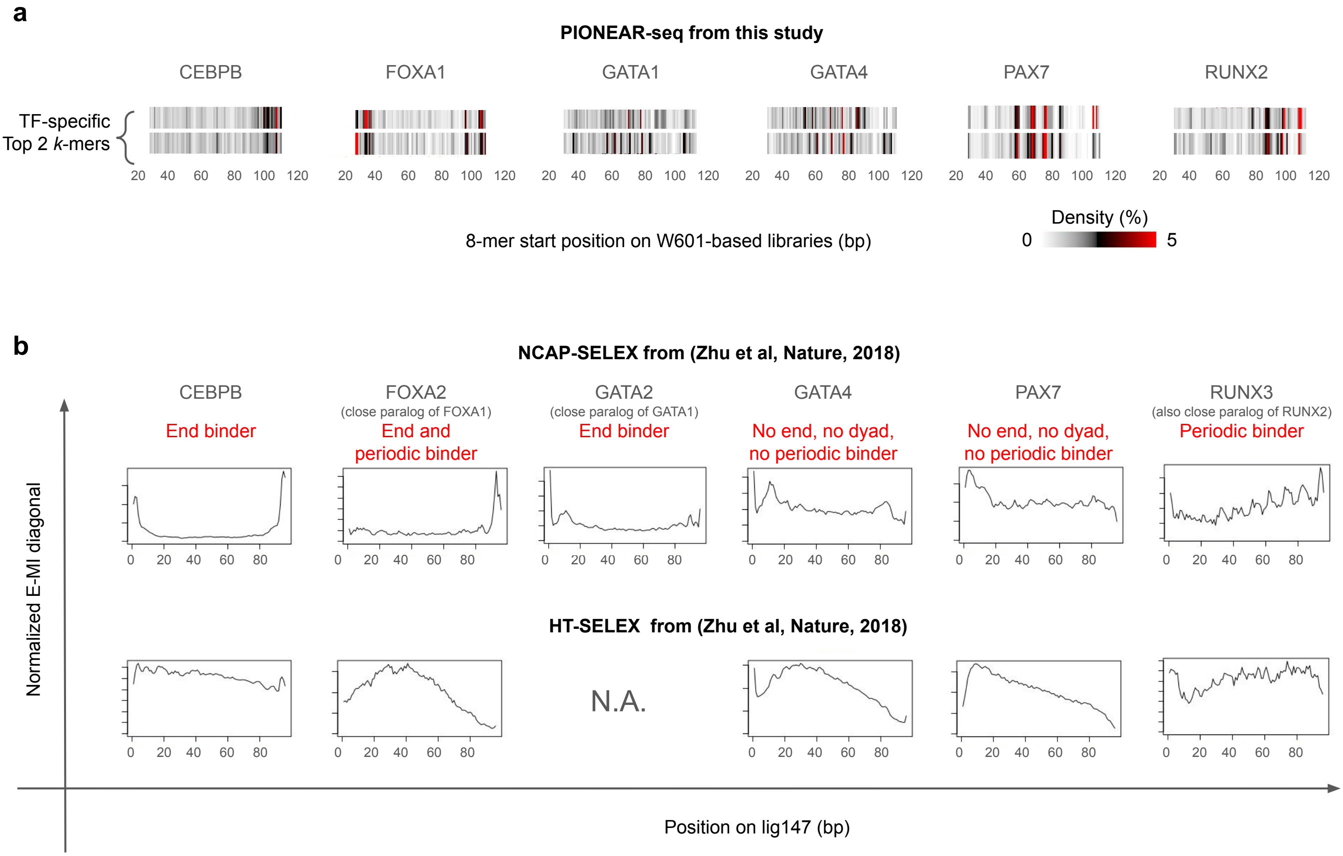


**Extended Data Fig. 8**

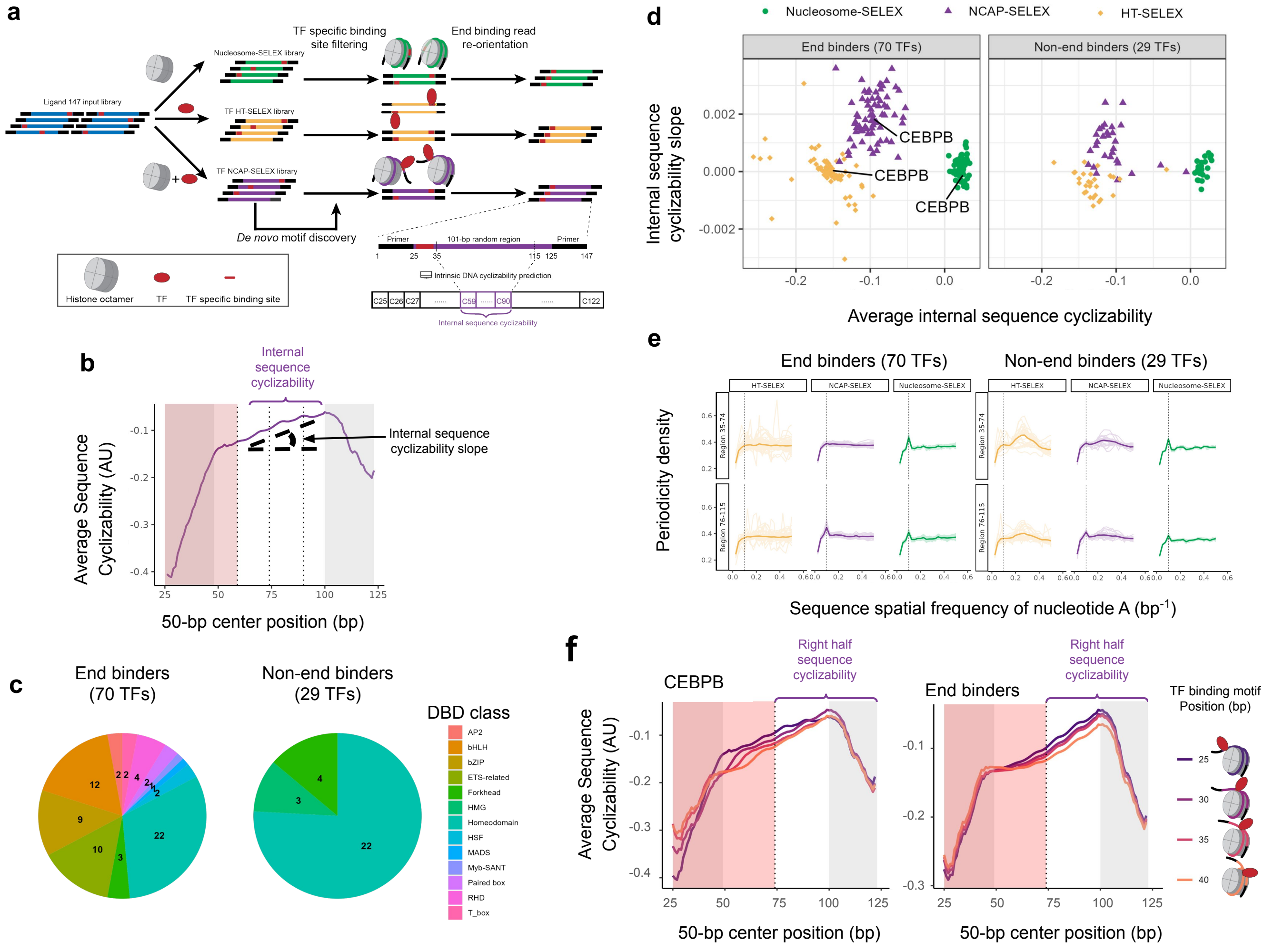


**Extended Data Fig. 9**

| HUGO TF name | CIS-BP Reference PWM (ID and Logo) | **Free DNA** Top k-mers | **NCP** Top k-mers |
|---|---|---|---|
| ASCL1 | M04135 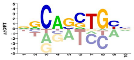 | **GCAGCTGC** | **AGCAGATG, GACAGCTG, AAAAAAAA** |
| CEBPB | M02851 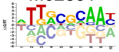 | **TTGCGCAA** | **TTGCGCAA** |
| FOXA1 | M04819 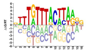 | **TGTTTACTT** | **TGTTTACTT** |
| GATA1 | M02738 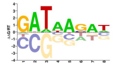 | **GATAAGAT** | **GATAAGAT** |
| GATA4 | M03066 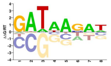 | **AGATAAGG** | **AGATAAGA, ATCAAGAT** |
| KLF4 | M04463 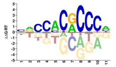 | **(G/A)CCACGCCC** | **TGCATAATT, ATTAGCAT , TTTGTTTAC, AAATTTTT** |
| OCT4 (*POU5F1* gene) | M05501 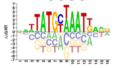 | **TATGCTAATT, TGCATAATTA,** | **TATGCTAATT** |
| PAX7 | M03292 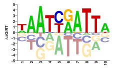 | **AATCGATT** | **AATCGATT** |
| PU.1 (*SPI1* gene) | M02965 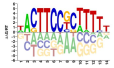 | **ACTTCCTCT** | **CTTCCTCTT** |
| RUNX2 | M03470 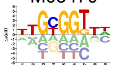 | **TTGCGGTT** | **TTGCGGTT** |
| ZNF143 | M02899 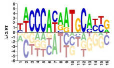 | **CCACAATGC** | **CCAGAATGC** |

**Extended Data Table 1: DNA Binding specificity for the TFs assayed by PIONEAR-seq**
