## Supplementary Information for "DNA flexibility regulates transcription factor binding to nucleosomes"

**Table of contents:**

1. **Methods**
2. **Supplementary Discussion**
3. **Extended Data Figure legends**
4. **Supplementary References**

### 1 - Methods

#### 1.1 Histone octamer assembly and purification

Purified recombinant *H. sapiens* canonical histones (H2A, H2B, H3.1 and H4) were purchased from Histone Source (Catalog SKU #: HH2A, HH2B, HH3 and HH4), aliquoted, lyophilized, and stored at -20°C prior to use. The lyophilized histones were resuspended in unfolding buffer (6 M guanidine hydrochloride, 5 mM DTT and 20 mM Tris-Cl, pH 7.5) to a concentration of 2-3 mg/ml. H2A, H2B, H3.1, and H4 were then combined at a molar ratio of 1.1:1.1:1:1. The sample was incubated at room temperature for 30 minutes before it was transferred into a 10K MWCO Slide-A-Lyzer osmotic cassette (ThermoFisher cat. #66380) and dialyzed for at least 16 hours at 4°C against 1 L refolding buffer (10 mM Tris 7.5, 2.0 M NaCl, 1 mM EDTA and 5 mM 2-mercaptoethanol). After dialysis, the sample was further dialyzed for 2 hours at 4°C in 1 L fresh refolding buffer and purified by size exclusion chromatography using a Superdex 200 Increase 10/300 GL column (Cytiva) (**Extended Data Fig. 1a)**. The FPLC fractions containing the octamer were pooled, concentrated to about 1.5 mg/ml using 10K MWCO Amicon Ultra spin columns (EMD Millipore) and stored at 4°C. In subsequent assays that required nucleosome assembly (PIONEAR-seq, EMSA, MNase-seq), histone octamers were always handled in the cold room (4°C) and within 1 month from their purification by size exclusion chromatography.

#### 1.2 Nucleosome Assembly

We assembled nucleosomes by slow, cold salt dialysis, essentially as described previously[1]. Briefly, DNA of interest (147-bp DNA produced by PCR) was quantified by $A_{260}$ and mixed in 50 µl of cold High Salt Buffer (10 mM Tris-HCl pH 7.5, 2.0 M KCl, 1 mM EDTA and 1 mM DTT). Typically, 2-10 µg of DNA were assessed with different amounts of histone octamer to find the optimal DNA:octamer molar ratio that produced the best yield of Nucleosome Core Particles (*NCPs)*, and which typically ranged between 1:1.2 and 1:1.8 (**Extended Data Fig.**

**1b**). The formation of NCPs was induced by placing each DNA:octamer sample in a distinct 7K MWCO Slide-A-Lyzer MINI dialysis device osmotic cap (#69562 ThermoScientific), which was slowly (>16 hours) dialyzed in the cold room from High Salt Buffer (see above) to Low Salt Buffer (10 mM Tris-HCl pH 7.5, 0.2 M KCl, 1 mM EDTA and 1 mM DTT) in a linear gradient using a peristaltic pump, as previously described[1]. The osmotic caps were then placed into 400 ml of Nucleosome Short-term Storage Buffer (10 mM Tris-HCl pH 7.5, 1 mM EDTA, 1 mM DTT) containing a stirring bar for at least 1 hour in the cold room. For each sample, the nucleosome assembly was assessed by Electrophoretic Mobility Shift Assay (EMSA) on commercial native NOVEX PAGE TBE gels (Invitrogen, EC62252/EC6365) stained by GelRed or ethidium bromide and imaged by Gel Doc EZ (Bio-Rad) or Odyssey (Li-COR) (**Extended Data Fig. 1b,c**).

#### 1.3 PIONEAR-seq DNA library design and production

##### 1.3.1 Templates

The PIONEAR-seq library design was based on four 147-bp DNAs (**Supplementary Table 1:** "PIONEAR-seq DNA sequence design"): 3 templates (*ALBN1, CX3CR1, NRCAM*) were of genomic origin and known to position nucleosomes *in vivo*[2]; 1 template, Widom 601 (W601), was of synthetic origin and was selected to have high affinity for histone octamers[3]. For each template, a pair of forward and reverse primers with a melting temperature of approximately 65°C (online ThermoFisher Tm calculator) were used in subsequent PCR amplification steps (primer sequences provided in **Supplementary Table 1**). Each primer pair was tested by PCR to be >99% specific to its own template as compared to the other templates. The 4 primer pairs were also tested to have a similar amplification efficiency for their cognate templates.

#### 1.3.2 Random tiles

In the design of the PIONEAR-seq input libraries, 20 consecutive bases in the internal portion of the template between the forward and reverse primers were modified to be random sequences ('random tiles'), so that each element of the library contained just one tile and the offset between tiles in different library elements was a multiple of 3 bp (**Supplementary Table 2:** "PIONEAR-seq Input Libraries").

#### 1.3.3 DNA Input libraries

For each template, ~30 tile-specific sublibraries of single-stranded oligos containing 20-bp random tiles at a certain position (*$N_{20}$ sublibraries*) were purchased as Ultramer DNA oligos (IDT DNA) and pooled together to be equimolar, thus creating 4 template-specific oligo sublibraries (*template sublibraries*). Each template sublibrary was separately amplified by PCR (KAPA HiFi Hotstart ReadyMix, 07958935001, KK2602, Roche) using its specific constant primers ("PIONEAR-seq DNA sequence design" in **Supplementary Table 1**) and an annealing temperature of 62°C. The PCR products were cleaned by Zymo-5 or Zymo-25 Clean&Concentrator spin column (Zymo Research). The 4 resulting DNA template sublibraries were assayed by D1000 TapeStation (Agilent) for the presence of a single ~147-bp DNA band and by NanoDrop to quantify the DNA yield, and then pooled together in an equimolar ratio to form the "DNA Input library" for PIONEAR-seq experiments.

### 1.4 MNase-seq assay and analysis

#### 1.4.1 MNase digestion

Nucleosomes for either single 147-bp DNA sequences (**Extended Data Fig. 1d, Supplementary Table 3:** "MNase-seq single sequences") or for the DNA input library (**Fig.**

**3b, Extended Data Fig. 1e,f, Extended Data Fig. 6a**) were reconstituted as described above (Methods section 1.2: “Nucleosome assembly”). Nucleosome assembly was assessed by EMSA to ensure the samples exhibited a weak band (<10% of total DNA intensity) at ~150 bp (residual free DNA) as compared to a strong band (>90% of total DNA intensity) at ~450 bp (NCP band). The MNase digestion was performed using a commercial MNase reaction kit (M0247S, NEB). The 2000 units/µl MNase was prediluted (1:100, 20 units/µl) in the manufacturer’s storage buffer from the kit and stored at -20°C for multiple uses. The Reaction Master Mix (RMM) was prepared fresh by mixing at a 10:1 ratio the 10x MNase Reaction Buffer and the 20 mg/ml (100x) BSA from the kit. The MNase reaction was prepared in a 1.5-ml tube on ice to a final volume of 40 µl by mixing 27 µl nucleosome sample, 6 µl RMM and 7.5 µl prediluted MNase, carried out in a water bath at 37°C for 10 min and blocked by adding 40 µl stop buffer (40 mM EDTA, 40 mM EGTA, 1% SDS, 1.5 mg/ml proteinase K). The reaction was checked by loading aliquots (4-8 µl) of NCP samples before and after MNase digestion on an EMSA TBE gel (Invitrogen) to ensure that the digested samples exhibited only the NCP band at ~450 bp and minimal band intensity for free DNA.

**1.4.2 Sample preparation for sequencing**

The blocked samples in stop buffer were incubated overnight at 37°C, and the digested DNA fragments were recovered with Zymo-5 Clean&Concentrator spin columns (Zymo Research), checked by D1000 TapeStation (Agilent) and prepared for sequencing using an NEBNext® Ultra™ DNA Library Prep Kit for Illumina® (E7370L, NEB) according to the manufacturer. Multiple prepared libraries were pooled together and sent to Azenta (previously called Genewiz) for Miseq and HiSeq 4000 sequencing (150 bp, paired-end) and demultiplexing. The forward (labeled ‘R1’) and reverse complement (labeled ‘R2’) 150-bp paired-end sequencing Fastq reads were merged with PEAR using default settings[4].

**1.4.3 Analysis of MNase cuts on single sequence NCPs**

Reads whose lengths were between 80 and 147 bp were aligned back to their template of origin (**Supplementary Table 3**) with the R function pairwiseAlignment (default setting except: type='overlap') and dismissed if there were any gaps or more than 5 mismatches, and if the alignment was less than 80 bp. Since the reads were filtered to be longer than 80 bases, the 5' end of the fragments always occurred at position between 1 and 67 bases, and the 3' end occurred at position between 81 and 147 bases, within the 147-bp DNA. The read counts of the cut position at the 5' end (at position between bases 2 to 67) and 3' end (at position between bases 81 to 146) were used separately to draw the left and right side of the distribution shown in **(Extended Data Fig. 1d).**

**1.4.4 Analysis of MNase cuts on PIONEAR-seq NCP input libraries**

To remove PCR duplicates, datasets of 'unique reads' were created by excluding duplicated reads, so that filtered reads with both identical tiles and identical end cuts were considered just once. Unique reads whose length was between 80 and 147 bp were assigned to the template (W601, *ALBN1, CX3CR1* or *NRCAM*) that exhibited the best alignment. Reads with no mismatch to the associated template were dismissed, since they represented fragments where the $N_{20}$ tile had been cut out and its position could not be identified. The remaining reads formed the "*QCed reads*" that then the algorithm scanned to identify the position of the $N_{20}$ tile within the template according to the original sublibraries **(Supplementary Table 2:** "PIONEAR-seq Input Libraries"**)**. For that, two groups of reads were selected. The first group represented fragments where the MNase cut happened (if any) on the left and right of the $N_{20}$ tile and included reads containing mismatches within an internal region no longer than 20-bp, which informed on the position of the $N_{20}$ tile in the reads. The second group represented fragments where the MNase cut happened within the $N_{20}$ tile and included reads that at one end contained up to 20 mismatches, which were also within the $N_{20}$ tile. These two read groups were merged, and then sorted according to their $N_{20}$ sublibrary of origin (**Supplementary Table 2**). To visualize whether the position of the $N_{20}$ tile modulates the accessibility of the nucleosomes, the MNase-digested $N_{20}$ sublibraries of the same template were visually compared as normalized distributions (%) of the MNase cuts on the left and on the right of the $N_{20}$ tile **(Extended Data Fig. 1e,f**). Finally, the QCed reads of different templates were compared and their percentages in the original pool of 4 template libraries were calculated, which quantified the relative degree of protection from MNase digestion between libraries based on distinct templates (‘After digestion’ in **Extended Data Fig. 6a**). Two independent replicates showed that the fragments from the W601-based library were the most abundant, indicating that the W601 template was the most protective compared to the three genomic templates (**Fig. 3b**). As a control, the same analysis performed on NCP input libraries not incubated with MNase (‘Before digestion’ in **Extended Data Fig. 6a**) revealed a very similar number of reads across different templates (~25% each), as expected since the DNA input library came from an equimolar pool of the 4 template sublibraries (see above, “1.3 PIONEAR-seq DNA library design and production”).

#### 1.5 GST-tagged TF expression for PIONEAR-seq

Gateway-compatible codon-optimized ORF sequences for the principal isoforms of the 11 human full-length transcription factor (TF) genes to express ASCL1, CEBPB, FOXA1, GATA1, GATA4, KLF4, OCT4 (gene *POU5F1*), PU.1 (gene *SPI1*), PAX7, RUNX2, and ZNF143 (**Supplementary Table 4:** “DNA sequences for TF expression”) were synthesized as gBlocks by Integrated DNA Technologies. They were then transferred by Gateway recombinational cloning into pDEST15 (ThermoFisher cat. #11802014) for expression as N-terminal GST fusion proteins and sequence-verified by Sanger sequencing. TF proteins were expressed by the PURExpress coupled *in vitro* transcription and translation kit (NEB) according to the manufacturer’s protocols at 30°C for 4 hours. Protein expression, size and concentration were determined by Western blot using a recombinant GST dilution series (Sigma) and anti-GST antibody (Abcam ab92; SIGMA SAB4301139) (**Extended Data Fig. 2a**), essentially as described previously[5]. Protein expression was repeated for each PIONEAR-seq selection round and stored at 4°C up to 48 hours prior to use.

#### 1.6 PIONEAR-seq assay

##### 1.6.1 NCP input library selection by TFs

The NCP input library was generated by assembling nucleosomes with DNA Input library. To capture TF:NCP complexes, each GST-tagged TF at a concentration of 50 nM was separately incubated with 200 µg of the NCP input library in 300 µl "PIONEAR-seq Binding Buffer" (10 mM Tris-HCl pH 7.5, 50 mM KCl, 5 mM NaCl, 1 mM MgCl, 3uM ZnSO4, 0.5 mM EDTA, 4% glycerol) supplemented with 1 mM DTT, 150 ng poly dI/dC and 40 µl pre-equilibrated Pierce™ Glutathione Agarose beads (ThermoFisher cat. #16101) ("NCP" conditions). As a negative control for the effect of histone octamer on TF:DNA specific interaction, the PIONEAR-seq assay was also performed on DNA input libraries not assembled on nucleosomes ("Free DNA" conditions). As a control for the nonspecific binding of the DNA and NCP samples to the glutathione beads, the input libraries were also incubated with Pierce™ GST peptide (PI20237 ThermoScientific) ("GST" controls). Each sample was incubated at 4°C in continuous rotation for >8 hours, then the glutathione agarose beads were collected by centrifugation (30 sec at 1000 rpm) and then washed 10 times with 200 µl cold PIONEAR-seq Binding Buffer. To amplify the pulled-down DNA, the beads were resuspended in 150 µl KAPA mix and 4 µl of an equimolar combination of the 4 template-associated primer pairs (**Supplementary Table 1:** "PIONEAR-seq DNA sequence design"). The PCR mix (180 µl) was divided into 4 PCR tubes and amplified for 10-12 PCR cycles with an annealing temperature of 62°C. The resulting PCR product was purified with Zymo-5 Clean&Concentrator spin column (Zymo Research), eluted in 20 µl water, assessed by D1000 TapeStation (Agilent) to contain a single band at ~147 bp and quantified by NanoDrop (*Round 1 libraries*). If further bands were present and the ~147-bp band did not encompass at least 95% of the total D1000 DNA signal, the 147-bp DNA band was separated by electrophoresis on a 3% TAE gel stained with ethidium bromide and extracted and purified with a MinElute Gel Extraction kit (Qiagen cat. #28604). In case the amount of DNA was insufficient for the nucleosome assembly (<2 µg), the DNA was further PCR amplified to obtain 4 µg.

#### 1.6.2 Rounds 2-4

During each of the subsequent 3 rounds of PIONEAR-seq selection, libraries from the previous round were (1) assembled into NCPs (or left as free DNA), then (2) incubated with the same TF (or mock GST), and then (3) PCR amplified, as described for Round 1. In the case of free DNA, a yield of 200 ng was sufficient to perform the next round. In the case of NCP libraries, a minimal yield of 2 µg was required for nucleosome assembly and libraries were PCR amplified using 160 µl PCR reaction mix (primer and KAPA mix) for a number of PCR cycles calculated to produce at least 4 µg DNA (*i.e.*, 4 cycles if starting with 250 ng), which was then purified with a Zymo-25 Clean&Concentrator spin column (Zymo Research), eluted in 40 µl $H_2O$ and checked by D1000 TapeStation (Agilent) for the presence of a single ~147-bp DNA band. The resulting PCR product was assessed by D1000 TapeStation (Agilent) and quantified by NanoDrop as described for Round 1.

#### 1.6.3 Sequencing library preparation

For the 11 TFs and the GST control, Round 4 NCP libraries and Rounds 1, 2, 3 and 4 Free DNA libraries were PCR amplified separately using primer pairs to isolate each template-specific library (*ALBN1, CX3CR1, NRCAM* or W601). Then, each template-specific library was PCR amplified (6-8 PCR cycles) twice (**Supplementary Table 1:** “PIONEAR-seq DNA sequence design”) as follows: (1) first with primers that flanked the library elements with Illumina Adaptors, including Unique Molecular Identifiers (UMIs) in the form of 6 random bases at the 3’ end of the template to favor the detection of sequencing clusters; (2) then with primers that flanked the Illumina Adaptors with multiplexing TruSeq dual-barcodes (5'-barcodes: i501-516, 3'-barcodes: i701-724) and P5/P7 promoters for Illumina sequencing. The barcoded libraries were quantified using a Qubit 4 Fluorometer (Invitrogen), inspected by D1000 TapeStation for the presence of a unique DNA band at ~300 bp, and pooled together to obtain ~3M reads for each TF library and ~15M reads for the GST and Input libraries. The libraries were then sent to Azenta (previously called Genewiz) for HiSeq 4000 sequencing (150 bp, paired-end) and demultiplexing.

### 1.7 PIONEAR-seq Sequencing Data Processing

#### 1.7.1 Tile Information

The forward (labeled 'R1') and reverse complement (labeled 'R2') 150-bp paired-end sequencing Fastq reads were merged with PEAR using default settings[4]. To remove PCR duplicates, datasets of 'unique reads' were created by excluding redundant reads, so that reads with both identical tiles (same DNA sequence and same position within the template) and identical UMIs were considered just once. The unique reads were trimmed at the 3' end to remove the 6-bp UMIs. Since PIONEAR-seq libraries by design are 147-bp long, trimmed reads with different lengths were removed. To extract the DNA sequence and the location of the tiles, the reads were aligned to the associated template (**Supplementary Table 1: "PIONEAR-seq DNA sequence design"**) by the R function pairwiseAlignment (default setting except: type='overlap', gapOpening=1000, gapExtension=0). This function furnishes an alignment score and the position of the base mismatches for each read. Given the template is 147-bp long and the random tile is 20-bp long, reads with an alignment score smaller than 127 were considered non-aligned and removed. The position of the first and last mismatch were compared to assign each read to its sublibrary of origin. For statistical power, datasets with less than 100K unique tiles were dismissed from subsequent analyses (*QCed reads*).

#### 1.8 Analysis of *k*-mer enrichment in PIONEAR-seq libraries

By definition, *k*-mers are substrings of length *k* within a sequence of nucleotides of length at least *k*. In the following analysis, *k*-mers refers to substrings found within the internal 16 bp of the $N_{20}$ tiles and evaluated for their role as specific DNA binding sites of TFs. Since the local composition of the template flanking the $N_{20}$ tiles could form TF binding motifs due to junction effects and bias the motif enrichment, 2 bp on both the 5' and the 3' end of the $N_{20}$ tiles were trimmed so that in the following analyses, *k*-mers refers to substrings found within the internal 16-bp sequences of the $N_{20}$ tiles (*16-bp* $N_{20}$) and evaluated for their role as specific DNA binding sites of TFs. Since the number of bp of the TF:NCP binding site that contributes to the binding specificity is not known *a priori* and depends on the TF and the DNA state (*i.e.*, free DNA or NCP condition), the QCed reads were analyzed using the SELEX package[6], which is based on a *de novo* DNA motif discovery approach that compares the most frequent *k*-mers present in a given foreground set of DNA sequences to the frequency of the same *k*-mers in a given background set. In the following analysis, the 16-bp $N_{20}$ of QCed reads of each TF (or GST) selected library (either as NCP or free DNA) were separately used as the foreground set against the background set formed by the input library based on the same template. To increase the computational efficiency of the de novo search, (1) datasets of unique tiles were created, so that 16-bp sequences coming from identical $N_{20}$ tiles at the same position within the template were considered just once, (2) for unique tiles datasets larger than 1M in the case of TFs and GST ( or larger than 5M in the case of input libraries), downsized datasets were created by random resampling 1M unique reads (or 5M in the case of input libraries) (*16bp downsized datasets*). The 16-bp downsized datasets of the input library were first evenly divided into training and test datasets for constructing hidden Markov models (HMMs) of different orders using the R package 'Selex'[6,7], which automatically selected the HMM with the highest $R^2$ between the observed *k*-mer probability vs. the predicted *k*-mer probability.

The enrichment method evaluated each *k*-mer length ranging from 5 to 15 bp, independent of any prior knowledge of the optimal motif length. For each *k*-mer length, the SELEX package created a table containing all the *k*-mers that are both enriched and present more than 100 times in the foreground set. For each *k*-mer, the following three variables were collected:

- Observed Count*:* the number of matches of the *k*-mer in the foreground set
- Expected Count*:* the number of matches of the *k*-mer in the background set, normalized by the relative ratio of the sizes of the background and the foreground sets.
- Enrichment*:* the ratio between Observed Count and Expected Count of the *k*-mer (**Fig. 1b,c,f, Fig. 2a, Extended Data Fig. 2b, Extended Data Fig. 3a-c** ). The increase of the *k*-mer quantifies the *k*-mer enrichment in the library upon selection.

For each *k*-mer length, the SELEX package also evaluated the Kullback-Leibler (KL) divergence between the *k*-mer frequency distribution in the selected library and the input library as a measure to quantify the gain in information content of the entire foreground set as compared to the background set. Namely, the KL divergence between a selected library and the input library is computed as follows:

$$KL(selected \,||\, input) \;=\; \sum_{s \in S_{100}} P_{selected}(s) \log \frac{P_{selected}(s)}{P_{input}(s)},$$

where $S_{100}$ is the set of *k*-mers appearing at least 100 times in the libraries, $P_{selected}(s)$ is the probability of the *k*-mer $s$ appearing in the selected library, and $P_{input}(s)$ is the probability of $s$ appearing in the input library[6]. As previously suggested, the optimal *k*-mer length of the sample was set as the one associated with the maximum value of the KL divergence[6]. To find the optimal *k*-mer length of a TF, the information content at different *k*-mer lengths across its libraries was manually inspected, which indicated 10-mers for OCT4, 9-mers for FOX1, KLF4, PU.1, ZNF143 and 8-mers for the remaining TFs (including the GST-mock control samples). With these lengths, a list of consensus *k*-mers was curated to identify one representative binding sequence *k*-mer for each library (**Supplementary Table 5:** "PIONEAR-seq *k*-mer Enrichment Summary"). These *k*-mers were visually compared with known PWMs from the Cis-BP database (**Extended Data Table 1**).

#### 1.9 Analysis of *k*-mer positional preferences in PIONEAR-seq libraries

##### 1.9.1 Pre-processed dataset resampling

In the pre-processed datasets, the number of unique reads with tiles at certain positions varies. While this variability could reflect a preference for the TF to bind certain regions of the nucleosome, it could also be due to PCR amplification biases that favor certain $N_{20}$ sublibraries, resulting in an apparent positional preference for certain *k*-mers at certain positions. To avoid this bias, each pre-processed dataset ("*QCed reads*" from **1.7**) was randomly resampled to create an 'isotiled' dataset that has the same number M of tiles at each position (Isotiled datasets*)*, where M is equal to the median value of the distribution of the $N_i$ tiles present at each position I. For the resampling of each $N_{20}$ sublibraries, the R command 'sample( $N_i$ , M, replace = ( $N_i$ < M))' was used.

##### 1.9.2 *K*-mer positional density profiles along the NCP template

The positional density profiles of a certain *k*-mer in the indicated library were evaluated within the 16-bp tiles of the associated isotiled dataset as follows. By convention, the position of a *k*-mer within a template was defined by the position of the *k*-mer's first base relative to the 147-bp template sequence, which was indexed with increasing numbers from the 5' to the 3' end. In the case of the W601-based libraries, which by design had 21-bp and 28-bp constant primer sequences at the 5' and 3' end, respectively, the position of 8-mers in the 16-bp tiles along the 147-bp template ranged from bp 24 (=1+21+2) to bp 110

(=1+147-28-2-8). Therefore, depending on the template and on the *k*-mer length, the *k*-mer positional density profiles exhibited slightly different ranges: 8-mers ranged from bp 23 to bp 112 in *ALBN1*, from bp 20 to bp 118 in *CX3CR1* and from bp 28 to bp 117 in *NRCAM*-based libraries. For the isotiled dataset, the matches of the *k*-mer within the 16-bp tiles were then used to generate a distribution of the *k*-mer positions relative to the 147-bp template sequence. Since different *k*-mers exhibited different numbers of matches, each *k*-mer distribution was normalized by the total number of matches to create its density profile, which allows one to compare the positional binding preferences between *k*-mers with different match counts (**Fig. 2b, Fig. 3a, Extended Data Fig. 3d, Extended Data Fig. 7, Extended Data Fig. 8a**).

### 1.10 CEBPB:NCP binding reaction followed by EMSA

#### 1.10.1 CEBPB expression and purification

Codon-optimized open reading frames of human CEBPB DNA binding domain (residues 244-345) and full-length (residues 1-345) (**Supplementary Table 4:** "DNA sequences for TF expression") were subcloned into pET28b vector between an N-terminal $His_{10}$-SUMO tag and a C-terminal eGFP tag. Recombinant CEBPB proteins were expressed in 330 mL LB cultures of LOBSTR *E. coli* cells[8]. Cells were cultured at 37°C to 0.5 $A_{600}$ and induced with 1 mM IPTG for 2 hours at 37°C. Cells were harvested by centrifugation at 2,700 g for 10 min, and the cell pellets were resuspended in 20 mL lysis buffer (50 mM HEPES at pH 7.5, 200 mM NaCl, 0.5 mM TCEP) supplemented with Complete protease inhibitor cocktail (Sigma), and mechanically lysed with 2 rounds of French Press, followed by centrifugation (20,000 g rpm for 30 min at 4°C). The supernatant was incubated with 0.33 mL of Ni-NTA beads (MCLAB) for 30 minutes at 4°C. The beads were then poured onto an Econo column (Bio-Rad). The flow-through was collected and was supplemented with concentrated NaCl and imidazole up to 1 M NaCl and 20 mM imidazole.  This flow-through was incubated with fresh 0.33 mL Ni-NTA beads (MCLAB) for 10 minutes at 4°C to capture the His-tagged CEBPB proteins and the resin was washed with 5 mL high-salt lysis buffer (50 mM HEPES at pH 7.5, 1 M NaCl, 20 mM imidazole, 0.5 mM TCEP) and then with 20 mL lysis buffer (50 mM HEPES at pH 7.5, 200 mM NaCl, 0.5 mM TCEP). CEBPB proteins were eluted from the beads by adding 4 mL of the elution buffer (lysis buffer with 400 mM imidazole). CEBPB samples in the elution were inspected and quantified by SDS-PAGE, Coomassie staining, and gel-based densitometry using BSA standards. Purified CEBPB samples were supplemented with filtered glycerol (final concentration = 10% glycerol), aliquoted (50-100 µl), flash-frozen, and stored at -80°C.

#### 1.10.2 DNA production and purification

Both ultramer oligos for 147 bp template sequences (**Supplementary Table 7:** "CEBPB-bound NCP EMSA single sequences") and oligos for the associated primers were purchased (IDT DNA). To produce ~100 µg of DNA of a specific sequence by PCR, a PCR Master Mix was prepared by thawing 2.5 ml of Taq 2× Master Mix (NEB, M0270L) on ice and adding 2.5 ml cold pure water. The template and the primers were added to the PCR Master Mix to a final concentration of 20 nM template Ultramer oligo and 400 nM each primer oligo. The reaction mixture was distributed in the 96 PCR tubes of a single plate placed on ice. PCR was carried out for 25 cycles after initial denaturation at 94°C for 30 s. For each PCR cycle, after 94°C heating for 30 s, the reaction was left to anneal at 58°C for 30 s, followed by 68°C elongation for 30 s, and a final 68°C elongation step for 5 minutes. After PCR, all 96 tubes were combined into one pool and diluted with Ion Exchange (IEX) buffer (10 mM Tris pH 7.8, 1 mM EDTA) to a final volume of 50 ml. To purify the DNA, the solution was applied to 2 ml of Q Sepharose Fast Flow ion exchange column (Cytiva 17051001) binding twice. The column was washed 3 times with 10 ml Low-Salt IEX buffer (IEX buffer with 0.3 M NaCl) to remove free mononucleotides and oligonucleotides, followed by elution with 10 ml high-salt IEX buffer (IEX buffer with 1 M NaCl) and by buffer exchange back to IEX buffer. The purified DNA was concentrated to < 0.2 ml using 10 kDa MWCO Amicon Ultra centrifugal filters (EMD Millipore), quantified by Nanodrop, checked by D1000 tape station (Agilent) for the presence of a single peak at ~147bp and stored at 4°C for usage within 1 month.

#### 1.10.3 CEBPB:NCP binding reaction followed by EMSA

For each nucleosome assembly reaction, 10 µg of DNA of a certain template were reconstituted as described above (Methods section 1.2). The binding of purified CEBPB (Methods section 1.10), as either the DNA binding domain or full-length protein, to individual DNA sequences, as either free DNA or DNA assembled into NCPs, was determined by multiple binding reactions (constant amount of DNA and variable amounts of CEBPB) followed by EMSA. All the reactions, including preparation and gel running, were performed in the cold room. Each DNA-binding reaction was carried out in 10 µl, of which (i) 5 µl of NCP (or free DNA) pre-diluted to 200 ng/µl (quantified by NanoDrop for DNA) in Nucleosomes Short-term Storage Buffer (10 mM Tris-HCl pH 7.5, 1 mM EDTA, 1 mM DTT), and (ii) 5 µl purified CEBPB (different amounts) stored in elution buffer (50 mM HEPES pH 7.5, 400 mM imidazole, 0.5 mM TCEP, 10% glycerol) containing 0.04% Triton-X 100 and 0.4 mg/mL bovine serum albumin (BSA) (NEB). NCP (or free DNA) samples were diluted to 0.04 mg/ml in cold Nucleosomes Short-term Storage Buffer. Frozen CEBPB samples were thawed on iced and supplemented with Triton-X and BSA and used for serial dilutions (dilution factor = ½). Both NCP (or free DNA) and CEBPB samples were used within 30 min of being diluted. Each binding reaction was carried out in one PCR tube by mixing 5 µL CEBPB sample with 5 µL NCP (or free DNA) samples. The PCR tubes were capped and incubated overnight in the cold room. The next day, samples were gently loaded (no loading buffer added) on cold, prerun, non-denaturing polyacrylamide gels (18 well 5% Criterion™

TBE Polyacrylamide Gel (Bio-Rad 3450048)) and electrophoresed in the cold room for 3 hr at 60 V. Gels were stained with ethidium bromide and imaged by Gel Doc EZ (Bio-Rad) or Odyssey (Li-COR).

#### 1.11 *In silico* modeling of the CEBPB-nucleosome complex using AlphaFold3

The AlphaFold3 server (alphafoldserver.com)[9] was used to model the structure of the CEBPB-nucleosome complex (**Fig. 2c, Extended Data Fig. 4**). The unmodified W601 DNA template sequence and the full-length *Xenopus laevis* histone protein amino acid sequences were taken from a Cryo-EM structure (PDB: 7OHC)[10]. Various W601 DNA mutants with CEBPB-binding motifs were derived from PIONEAR-Seq results (**Supplementary Table 6:** "Sequences used for AlphaFold3 structure prediction"). The 8-mer CEBPB-cognate motif (TTGCGCAA) within the W601 DNA sequence was introduced at three different positions to evaluate CEBPB's site selectivity for nucleosome association: at bp 68 (SHL approximately -0.5), bp 65, and bp 71, with an additional 1 bp flanking DNA on both sides (ATTGCGCAAT). The CEBPB protein sequence, corresponding to residues Val259-Pro335, was taken from the crystal structure of the CEBPB-DNA complex (PDB: 8K8D)[11].

#### 1.12 *In silico* modeling of the CEBPB-nucleosome complex using Rosetta3

First, the nucleosome core particle containing *Xenopus laevis* histones H2A (Ala12-Lys119), H2B (Lys27-Lys125), H3 (Leu20-Ala135), and H4 (Lys16-Gly102) and a 146-bp palindromic Alpha-satellite sequence was selected as a modeling template (PDB: 1AOI)[12]. Using web 3DNA 2.0 (w3DNA 2.0)[13], the W601 template containing the CEBPB binding sequence was introduced.

The 146 bp palindromic Alpha-satellite sequence from the crystal structure (PDB: 1AOI) was:

5’-ATCAATATCCACCTGCAGATTCTACCAAAAGTGTATTTGGAAACTGCTCCATCAAAAGG
CATGTTCAGCTGAATTCAGCTGAACATGCCTTTTGATGGAGCAGTTTCCAAATACACTTTT
GGTAGAATCTGCAGGTGGATATTGAT-3’

This DNA was mutated to W601 sequence including the CEBPB binding sequence (TTGCGAAA) at bp 68 (SHL -0.5), derived from PIONEAR-Seq results (**Fig. 2b**):

5’-TCGAGAATCCCGGTGCCGAGGCCGCTCAATTGGTCGTAGACAGCTCTAGCACCGCTT
AAACGTGTA**TTGCGAAA**TATATTGGCGTTTTAACCGCCAAGGGGATTACTCCCTAGTCTC
CAGGCACCTGTCAGATATATAGATCCGAT-3’.

After removing metal and water molecules from the PDB structure, the structure was renumbered and cleaned using the Rosetta3 score_jd2 application with the following flags:

*score_jd2.static.linuxgccrelease -renumber_pdb -out:pdb -ignore_unrecognized_res -s 1aoi_mutated.pdb*

Then, using the Rosetta3 N-terminal acetylation application, each histone molecule in the nucleosome complex underwent N-terminal acetylation to avoid artifactual charge interactions caused by the N-termini of the truncated histones:

*N-terminal_acetylation.static.linuxgccrelease -s chainC.pdb -nstruct 1 -out:pdb*

The complex structure with the N-terminally acetylated histones was renumbered and cleaned again using score_jd2 with the following flags:

*score_jd2.static.linuxgccrelease -renumber_pdb -out:pdb -ignore_unrecognized_res -s starting_1aoi_mutated.pdb*

It then went through Rosetta FastRelax with the following flags: *relax.default.linuxgccrelease -s starting_1aoi_mutated_Nac.pdb -scorefile relax.sc -nstruct 10 -relax:fast -out:prefix relax_ -ex1 -ex2aro -dna_move*

The lowest score from the 10 relaxed structures was selected as a template nucleosome structure to generate a CEBPB-bound nucleosome structure for comparative modeling.

The relaxed nucleosome model was aligned with the existing crystal structure of the CEBPB consensus motif in complex with the C-terminal bZIP domain of CEBPB (Lys269-Pro337; PDB: 6MG1)[14]. Specifically, the TATTGC DNA sequence was used as an alignment anchor, which translocated the CEBPB-DNA complex near the SHL -0.5 position of the nucleosome, making the CEBPB dimer sit perpendicular to the globular histones.

Next, the CEBPB dimer and the nucleosome PDB files were merged, renumbered, and cleaned using score_jd2 application (score_jd2.static.linuxgccrelease -renumber_pdb -out:pdb -ignore_unrecognized_res -s 1aoi_CEBPB-6mg1_combined.pdb) and used as an initial template for comparative modeling. Rosetta FastRelax was used to generate 150 relaxed structures with the following flag:

*relax.default.linuxgccrelease -s 1aoi_CEBPB-6mg1_combined_cleaned.pdb -scorefile relax.sc -nstruct 150 -relax:fast -out:prefix relax_ -ex1 -ex2aro -dna_move*.

The lowest energy model was used as a representative structure (**Fig. 2d**).

#### 1.13 Cyclizability prediction of DNA sequences

DNA sequence cyclizability represents the ability of a sequence to form a DNA loop and has been measured experimentally in a high-throughput manner by loop-seq technology for 50-bp random libraries inserted into 100-bp linear DNA[15]. By using these loop-seq data, two machine learning tools have been created to predict the cyclizability of any given 50-mer (see below). In this study, the sequence cyclizability was used as a proxy for sequence flexibility and in the text we use these two properties of a DNA sequence interchangeably. Since the cyclizability has a resolution of 50 bp, to profile cyclizability of a 147-bp sequence (as either a single template (**Fig. 3e**), an element in the PIONEAR-seq libraries (**Extended Data Fig. 6c**) or NCAP-SELEX libraries (**Fig. 4b**)), we split the sequences into 98 50-mer sliding windows with the first window being the 50-mer centered on bp 25 of the template and the last window being centered on bp 122 (**Fig. 3c**). The cyclizability of each 50-mer is predicted by two different deep-learning methods: the first one[16] was developed by Taekjip Ha's lab (Harvard Medical School, Boston), who had developed the original loop-seq technology[15], and is used in analyses presented in **Figures 3c-e, 4b,d-f, 5a,b** and **Extended Data Figures 6c, 9b,d,f**. The method of Park *et al.*[16] provides deep-learning models for cyclizability prediction by training on denoised loop-seq datasets[15], resulting in accuracy comparable to experimental precision. Predictions of template and reverse complementary sequences highly correlate, indicating that the method can be applied to various settings regardless of the DNA strand analyzed. Results from this method were consistent with those obtained by an alternative deep learning approach, DNAcycP[17], which also predicts 50-mer DNA cyclizability (**Extended Data Fig. 6b)**. For both models, we followed default instructions by the developers for installation and running predictions for the 50-mer cyclizability.

#### 1.14 Sequence cyclizability in NCAP-SELEX screen data

##### 1.14.1 NCAP-SELEX screen data filtering and motif enrichment

The FASTQ files of published ligand 147 DNA sequence libraries, including the input cycle, nucleosome-SELEX bound cycle-4, HT-SELEX cycle-4, and N(ucleosome)CAP-SELEX cycle-4 for 195 TFs[18] were downloaded from the European Nucleotide Archive under accession #PRJEB226841; we refer to these data collectively as "NCAP-SELEX screen data". Paired-end reads were merged using PEAR with default settings[4]. To QC the library reads, duplicated reads in each library were counted just once, while reads either with incorrect lengths of the 101-bp random region in the internal portion of the ligand or with ambiguous nucleotides (*i.e.*, N's) were filtered out. The QCed reads of the input library were then evenly divided into training and test datasets for constructing hidden Markov models (HMMs) using the R package 'Selex'[6,7], which automatically benchmarked HMMs of order 0 to 6 and selected the HMM of order 6 because it exhibited the highest $R^2$ (0.998) between the observed *k*-mer probability against the predicted *k*-mer probability. For each QCed HT-SELEX library, the selected HMM determines the ability of a TF to specifically bind on free DNA by calculating the information gain, defined as the KL divergence between the *k*-mer frequency distribution in the selected library and the input library (Methods section 1.8: "Analysis of *k*-mer enrichment in PIONEAR-seq libraries"). A threshold of 0.1 for the HT-SELEX information gain was then set to filter out TFs whose DNA binding specificity was weak and furnished 137 TFs with strong motif specificity for further analysis (**Supplementary Table 8:** “NCAP-SELEX TFs”). The QCed NCAP-SELEX libraries for these 137 TFs were used to discover their binding specificity as a collection of enriched 7-mers as compared to the input library. For that, the 6th-order Markov model already constructed by the ligand 147 input library using the R package ‘Selex’[6,7] found the most represented 7-mers (at least 100 counts) in each QCed NCAP-SELEX library, and calculated the enrichment of each 7-mer as the fold-change between its observed counts and its expected counts (expected counts are the ones found in an input library of the same size of the NCAP-SELEX library). For each TF, 7-mers with enrichment <1 were filtered out, and then the top 10 enriched 7-mers (along with their reverse complements) were considered to describe the TF’s binding specificity (*TF specific 7-mers*).

#### 1.14.2 End binder classification and end binding read filtering

For each TF, the positions of the TF specific 7-mers were scanned on QCed reads from the TF NCAP-SELEX, TF HT-SELEX, and Nucleosome-SELEX libraries. For each dataset, the end binding reads, defined as the reads with the specific 7-mers occurrences within 10-bp to either end of the 101-bp random region (*i.e.*, 25 - 34 bp for the 5’ end of 147-bp reads), were selected. To call end (and non-end) binding TFs, the percentage of end binding reads in the TF NCAP-SELEX reads was used: if greater than 8%, the TF was called an end binder (70 TFs), while if less than 6% it was called a non-end binder (29 TFs) (**Supplementary Table 8**). Information regarding the DNA binding domain classes of end and non-end binders was collected from TFClass [PMID: 25361979] (**Supplementary Table 8**). Then, the end binding reads of TF NCAP-SELEX, TF HT-SELEX, and Nucleosome-SELEX libraries were further filtered to ensure that the internal 81-bp region of the reads did not contain any further TF specific 7-mer occurrence. They were then re-oriented to have the end binding 7-mers always on the left side and aligned according to their expected dyad position. The forward and reverse primers from the NCAP-SELEX data screen design, which are the constant illumina sequencing adaptors[18], were added back to the ends of the reads to extend the re-oriented random regions up to 147 bp (*re-oriented end binding reads*).

#### 1.14.3 Profiles of average sequence cyclizability

The cyclizability of each re-oriented end binding read was predicted using the models by Park *et al.* (unpublished)[16] described above (Methods section 1.13: "Cyclizability prediction of DNA sequences"), which furnished multiple profiles (one per read) for each dataset. At each 50-bp center position (from bp 25 to bp 122), the profiles were averaged and the 95% confidence intervals were calculated by fitting a t-distribution with the mean and standard deviation derived from the cyclizability score distribution at the position. Namely, the confidence interval of the average cyclizability score at position $p$ bp is defined as follows:

$$CI_p = [\underline{x}_p - t_{n-1}(0.025)\times\frac{s_p}{\sqrt{n}}, \underline{x}_p + t_{n-1}(0.025)\times\frac{s_p}{\sqrt{n}}],$$

where $n$ is the number of re-oriented end-binding reads, $\underline{x}_p$ and $s_p$ are the mean and standard deviation of the cyclizability scores of the 50-bp centered at position $p$ bp. For each set of re-oriented end binding reads, the internal sequence cyclizability slope is calculated by fitting a linear regression to the profiles of average sequence cyclizability formed by the internal 81-bp region parsed in 32 50-mers (**Extended Data Fig. 9f**). The internal sequence cyclizability slope of re-oriented end binding reads from NCAP-SELEX libraries of end binders versus non-end binders was compared with a two-tailed Wilcoxon rank sum test (**Fig. 4e**).

#### 1.14.4 Nucleotide periodicity

For each set of re-oriented end binding reads, the internal 81-bp DNA sequence was separated into two halves: one from base 35 to 74 and the other from base 76 to 115 according to the the positional numbering system of the 147-bp reads. Each half was converted into a binary vector with 1 indicating the occurrence of nucleotide 'A' and 0 for the other three nucleotides. The binary vectors of each half were processed by Fast Fourier Transform to obtain the spectrum periodicity density of nucleotide ‘A’, which were later averaged across all reads in the library (**Fig. 4c**, **Extended Data Fig. 9e**). Similar results were obtained using other nucleotides and dinucleotides.

#### 1.14.5 Internal binding read filtering

For each end binder, QCed NCAP-SELEX reads were scanned for the presence of its binding sites at distinct internal positions within the 147-bp DNA sequence. Besides the set of reads that contained the TF specific 7-mers at 25 - 34 bp from the ends as described above, read sets with 7-mers found within 30 - 40, 35 - 45, or 40 - 50 bp from the nucleosome ends were generated. For each set, the reads were re-oriented to have the TF binding sites on the left side, profiled for cyclizability and averaged (**Extended Data Fig. 9f**). For each TF, the read sets were then compared for the average sequence cyclizability of the “right half sequence”, which spans from 51 to 147 bp and by design does not contain binding sites in any of the sets. The sliding windows for calculating the cyclizability of the right half sequences were centered from position 75 bp to 122 bp. The averaged cyclizability scores within the right half sequences were then compared across the sets grouped by distinct TF binding positions with a one-way repeated measures ANOVA test where subjects are TFs (**Fig. 4f**).

### 1.15 Cyclizability analysis of nucleosome sequences in K562 cells

#### 1.15.1 K562 nucleosome position calling

The bam file of a K562 MNase-seq experiment aligned to hg19 was downloaded from ENCODE[19] under the accession code ENCFF000VMJ. Nucleosome positions were called using DANPOS3 with default settings and peaks of length 140 bp were filtered for further analysis[20]. To correct for *k*-mer distribution bias in MNase-seq fragments, Seqoutbias was used with read-size 101, pdist 100:400, and default settings for the remaining parameters[21]. The called nucleosome positions from the pre-correction signal were verified to agree with the post-correction signal using Integrative Genomics Viewer (IGV). The called positions were then converted into coordinates in hg38 with liftover[22].

#### 1.15.2 Classification of CEBPB-bound/unbound nucleosomes

ChIP-seq peaks and irreproducible discovery rate (IDR)-thresholded peaks for CEBPB in K562 cells were downloaded from ENCODE[19] under the accession codes ENCFF309ZJM and ENCFF712ZNR. The CEBPB binding site motif and itspositions on genome hg38 were identified as ATTGCAYAAY by Homer using the IDR thresholded peaks[23] (**Supplementary Table 9**: “K562 nucleosomes with CEBPB binding sites”). Bedtools intersect was used to separate motif positions covered by IDR thresholded ChIP-seq peaks vs. not overlapping any ChIP-seq peaks. Afterwards, the nucleosome positions with a CEBPB motif match covered by IDR thresholded ChIP-seq peaks at the end 30 bp were identified as “CEBPB-bound nucleosomes” (**Supplementary Table 9**). As a negative control set of nucleosomes not bound by CEBPB (“CEBPB-unbound”), nucleosomes not overlapping any CEBPB ChIP-seq peak and containing a CEBPB motif at the ending 30-bp were used[24]. Nucleosome DNA sequences were extracted from the reference genome hg38 using Homer tools, reoriented to have the CEBPB binding motif match on the left side and profiled for DNA cyclizability by 50-bp sliding windows as described above (**Fig. 5b**).

### 2 - Supplementary Discussion

#### 2.1 Systematic comparison between PIONEAR-seq results from this study and NCAP-SELEX results from Zhu *et al., Nature* (2018)

To systematically compare our results with those obtained by Zhu *et al.*[18] using NCAP-SELEX technology, we re-analyzed the enriched-sequence-based mutual information (E-MI) diagonal signal of NCAP-SELEX data obtained using 200-mers containing 154-bp randomized regions ("lig200" libraries) and for the 147-mers containing 101-bp randomized regions ("lig147" libraries).

##### 2.1.1 Preferential binding to nucleosomal DNA

In Zhu *et al.,* the lig200 NCAP-SELEX E-MI diagonal signal was used to evaluate the ability of a TF to recognize nucleosomal DNA in the presence of free DNA flanking the nucleosome. By quantifying the extension of the E-MI diagonal signal towards the lig200 central region, which is expected to be protected by the nucleosome from TF penetration, Zhu *et al.* highlighted VSX1 as a nucleosomal DNA binder because its signal had a very short minimum region (~53 bp) and SREBF2 as a free DNA binder because of its long minimum region (~104 bp) (**Extended Data Fig. 2g)**. Similar inspection of the E-MI diagonal signal profiles for lig200 NCAP-SELEX data from *Zhu et al.* showed that TFs (or close paralogs) that bound nucleosomes according to our PIONEAR-seq data (**Extended Data Fig. 2e**) were characterized by protection profiles similar to nucleosomal DNA binders in the NCAP-SELEX data, such as VSX1, while TFs that showed information gain below our threshold for detecting nucleosome binding in our PIONEAR-seq NCP libraries (**Extended Data Fig. 2f)** behaved similarly to free DNA binders, such as SREBF2 (**Extended Data Fig. 2h**).

##### 2.1.2 Positional binding preferences of TFs within nucleosomes

Of the 7 factors that we found to bind nucleosomal DNA by PIONEAR-seq (**Fig. 1b**), Zhu *et al.* assessed 3 of them directly (CEBPB, GATA4, PAX7) and close paralogs for another 3 (RUNX3 for RUNX2, FOXA2 for FOXA1, GATA2 for GATA1) for binding to nucleosomes built on 147-mers containing 101-bp randomized regions ("lig147"), which they used to call TF-nucleosome binding modes (*e.g.*, end binding, dyad binding); OCT4 and its close paralog POU5F2 were not assayed by lig147 NCAP-SELEX in Zhu *et al*. Since the lig147 NCAP-SELEX libraries are based on synthetic sequences selected to have affinity for histone octamers, we compared them with our PIONEAR-seq libraries based on the synthetic nucleosome position sequence W601. For 4 of these 6 TFs (CEBPB, FOXA1/2, PAX7,RUNX2/3), our W601-based PIONEAR-Seq data and the lig147 NCAP-SELEX data were in partial agreement with each other (**Extended Data Fig. 8)**:

- RUNX3 was called a periodic binder by Zhu *et al.* (and as apparent in Fig. 3 of Zhu *et al., Nature*, 2018 and in **Extended Data Fig. 8b** in our manuscript), consistent with our finding of periodic binding by its paralog RUNX2 by PIONEAR-seq for W601-based nucleosomes (**Fig. 3a**).
- CEBPB was called an end binder by Zhu *et al.*, consistent with our finding by PIONEAR-Seq that CEBPB binds its optimal recognition site TTGCGCAA at or near one of the ends of W601-based nucleosomes (**Fig. 2b**). However, we also found that some sub-optimal binding sites (*e.g.*, TTGCGWAA and the reverse complement TTWCGCAA) were preferentially bound at the dyad (**Fig. 2b**), an observation which we validated by molecular modeling (**Fig. 2c,d**) and EMSAs (**Fig. 2e**). This switch to a preferential alternate binding mode of CEBPB, observed for sub-optimal sequences (here, a 1 nucleotide mismatch from CEBPB's optimal recognition 8-mer), was not reported by Zhu *et al.*.
- FOXA2 was called both an end binder and a periodic binder by Zhu *et al.*, despite end binding being far more pronounced in their data than periodic binding (and as apparent in Fig. 3 of Zhu *et al., Nature*, 2018 and in **Extended Data Fig. 8b** in our manuscript). Those data are consistent with our finding of strong preference for end

binding, with only modest periodic binding, for its paralog FOXA1 by PIONEAR-seq using W601-based nucleosomes (**Extended Data Fig. 7b and S8a**).

- Zhu *et al.* did not call a preferential binding mode for PAX7. However, in our re-analysis of their data, we found that it showed a preference for binding one end of nucleosomes and modest periodic binding to nucleosomes (as apparent in Fig. 3 of Zhu *et al., Nature*, 2018 and in **Extended Data Fig. 8b** in our manuscript), whereas we found PAX7 to bind at the dyad and in a periodic manner by PIONEAR-seq on W601-based nucleosomes (**Fig 3a)**.
- GATA2 was an end binder in Zhu *et al.*, whereas its paralog GATA1 showed dyad and off-dyad binding by PIONEAR-seq on W601-based nucleosomes (**Fig. 7c**).
- GATA4 showed modest near-end binding in Zhu *et al.* (as apparent in Fig. 3 of Zhu *et al.* and in **Extended Data Fig. 8b** in our manuscript). This was consistent with the near-end binding that we observed for GATA4 binding to ATCTTATC, although we also observed dyad and off-dyad binding by GATA4 to that *k*-mer; however, intriguingly, we instead observed dyad and off-dyad binding to other 8-mers, with minimal near-end binding (**Extended Data Fig. 7d**).

However, we observed larger differences when comparing the binding modes called by Zhu *et al.* with the binding modes obtained by PIONEAR-seq for nucleosomes assembled on genomic nucleosome positioning sequences (*CX3CR1, NRCAM, ALBN1*), instead of W601 sequence. For example, RUNX3 was called a periodic binder by Zhu *et al.*, consistent with our finding of periodic binding by its paralog RUNX2 by PIONEAR-seq for W601-based nucleosomes (**Fig. 3a**), but RUNX2 bound to the ends of *ALBN1* and *CX3CR1* nucleosomes and to end, dyad, and near dyad sites on *NRCAM* nucleosomes (**Fig. 3a**). These differences highlight that the nucleosomal sequence context regulates the positional preferences of TF binding to nucleosomes; thus, the pioneer binding mode inferred in a certain DNA context by a TF may not generalize to nucleosomes assembled on other sequences.

Another novel key finding that we report in our manuscript is that the binding mode of a TF is further dependent on the particular binding site sequence (**Fig. 2b**), rather than being a constant binding mode across all its recognition sites.

**2.2 Previous evidence for TFs failing to bind nucleosomes**

For 4 of the 11 TFs we assayed by PIONEAR-seq, we did not detect sequence-specific binding. The lack of nucleosome binding observed by PIONEAR for these 4 TFs (ZNF143, KLF4, ASCL1 and PU.1) was supported by previous evidence in the literature:

- For ZNF143, we found just one published study suggesting the possibility of *in vivo* nucleosome binding from ChIP-seq data[25]. However, this possibility comes from results that were subsequently criticized in a study from the De Wit lab[26] as being due to experimental artifacts from cross-reactivity of the ZNF143 antibody with CTCF. Quoting from that bioRxiv preprint: *"After systematically re-analysing the ZNF143 ChIP-seq data, we posit that the Proteintech anti-ZNF143 polyclonal antibody recognises CTCF. in addition to ZNF143. To our knowledge, the only studies that report an overlap between ZNF143425 and CTCF are the ones that utilized this antibody."*
- For KLF4, *in vitro* binding to nucleosomes containing a KLF binding site has been reported[27], but to a much lower degree than other pioneers (*i.e.*, OCT4) (*"Klf4 was able to bind LIN28B-nuc with a higher apparent $K_d$ value compared to free DNA, indicating substantial nucleosome binding, but at a lower affinity than to free DNA"*) and in a partially nonspecific manner ("*both specific and non-specific interactions contribute to [Klf4] binding to nucleosomes in vitro*" and "*[Klf4 protects] both specific and non-specific nucleotides on LIN28B-nuc, supporting the non-specific contribution [of Klf4 bonding] to nucleosomes as seen in EMSA competition experiments*"). Since those observations do not distinguish between specific and nonspecific binding to nucleosomes, they are not in conflict with our results that KLF4 sequence-specific binding to

nucleosomes was below the detection limit in our PIONEAR-seq data. Moreover, in a follow-up study, Zaret and coworkers again found that strong reprogrammers (*i.e.*, OCT4) exhibited higher nucleosome binding than KLF4 and other TFs with supporting roles in reprogramming (Garcia *et al., Mol. Cell* (2019) 75(5):921-932, PMID: 31303471), again in agreement with the low level of KLF4 pioneer binding detected by PIONEAR-seq.

- For ASCL1, which as a bHLH TF that requires dimerization to bind DNA, nucleosome binding for the homodimer was previously found not to be detectable at the nanomolar range and required heterodimerization with the bHLH interacting partner E12alpha to bind nucleosomes at nanomolar affinity[2].
- For PU.1, an ETS factor which was shown to bind nucleosomes with nanomolar affinity[2], subsequent EMSA experiments on W601 nucleosomes tiled with PU.1 binding sites suggested that nucleosomes are a barrier to PU.1 binding and that sites near the nucleosome entry site (SHL +5.5, and strongly reduced binding at SHL +5) were at least partially accessible for binding by PU.1[28]. Notably, the nucleosomes used in the EMSA experiments that showed PU.1 binding with nanomolar affinity[2], were assembled on DNAs 160-162 bp long. A recent cryo-EM study found PU.1 to bind an ETS site at SHL +5.5 near the exit site of a *CX3CR1* nucleosome assembled on a 162-bp DNA and in their structures did not observe binding to GGAA core ETS sites at SHL -1.5 and -1.0[29]. Altogether, these studies suggest that PU.1 can bind near the entry/exit sites of nucleosomes, but is restricted from binding to internal sites on nucleosomes. In contrast to the experimental design of these prior studies, the NCPs we used for PIONEAR-Seq were assembled on 147-bp DNAs, rather than on ~160-162 bp DNAs. Moreover, the ends of our 147-bp DNAs were complementary to primers for use in PCR amplification of the libraries, with the 20-bp random regions tiling across the nucleosomes

from about SHL -4.5 to +4.5, where PU.1 might not have been able to bind its recognition sites.

#### 2.3 Comparison of PIONEAR-Seq to prior assays that have evaluated NCP:TF interactions

To our knowledge, our study is the first high-throughput study of the binding specificity of TFs to nucleosomes that combines the use of genomic sequences to assemble NCPs, insertions of random sequence, SELEX and sequencing. In particular, no prior study has explored weaker positioning sequences with such high-throughput approaches for detecting sequence-specific DNA binding preferences of TFs for nucleosomes. To substantiate this claim, we review here earlier studies that used some of these features to determine the DNA binding specificity of TFs to nucleosomal DNA:

- **Nucleosome positioning sequences of genomic origin**: Earlier studies have used weaker nucleosome positioning sequences of genomic origin, but not within a high-throughput binding site enrichment protocol like SELEX followed by sequencing. For example, in a study from the Zaret lab[2], which suggested the three genomic sequence templates that we used in this study (*ALBN1, CX3CR1, NRCAM*), the sequences were not mutated to assess the affinity of potential TF binding sites. Moreover, these studies used extended nucleosome positioning sequences longer than 147 bp, which is considered the optimal length to assemble NCPs[12].
- **W601-based sequences**: With the same purpose, other studies have applied high-throughput approaches that tile the W601-based nucleosomes with specific binding sites of the assayed TFs, most notably for OCT4 and SOX2[30] and P53[31]. However, these studies, which evaluated the positional preferences of TF binding to nucleosomes by Illumina sequencing, assessed binding to just one TF binding sequence across a W601-based nucleosome[30] or two TF binding sequences at just 7 positions across a W601-based

nucleosome[31], whereas by PIONEAR-seq we assayed for binding to essentially all possible TF binding sequences across the entire internal ~100-bp region of the nucleosomes.

- **Largely random nucleosome positioning sequences:** To reveal nucleosome binding and positional preferences for TFs, the NCAP-SELEX technology developed by the Taipale lab[18] assembled nucleosomes on 147-bp sequences that contained an internal 101-bp random region. These libraries were enriched by a few selection rounds for both histone octamer affinity and TF binding and then illumina sequenced, thus furnishing a high-throughout read-out of the TF nucleosome binding and positional preferences. However, in our re-analysis of these NCAP-SELEX data (**Fig. 4**), we found that the NCAP-SELEX enrichment cycles selected libraries that were more favorable for nucleosome formation (*i.e.*, more flexible and with greater nucleotide periodicity) than genomic nucleosome positioning sequences. This could create biases, since we found that the sequence context of nucleosomes influences the positional preferences of TF binding to nucleosomes (**Fig. 3a**).

### 3 - Extended Data Figure legends

#### Extended Data Fig. 1: Control experiments for NCP assembly in PIONEAR-seq libraries

**(a)** FPLC size-exclusion chromatograms of the *in vitro* assembled histone octamers. The fractions of histone octamers collected for the subsequent reconstitution of nucleosome core particles (NCPs) are highlighted (top segment). The chromatograms are representative of multiple replicates.

**(b)** EMSA gel results showing the *in vitro* reconstituted NCPs obtained from the indicated ratio of the FPLC-selected histone octamer (a) vs. the DNA input library, which came from an equimolar pool of the DNA libraries based on *ALBN1, CX3CR1, NRCAM* and W601 templates. Multiple octamer concentrations were assayed for the same DNA concentration (x-axis) to evaluate the NCP reconstitution optimal sample (label in bold) that exhibited minimal presence of residual free DNA. The optimal sample was used for TF binding and TF:NCP immunoprecipitation.

**(c)** Nucleosome assembly in subsequent rounds of selection. PIONEAR-seq DNA libraries selected by a certain TF were PCR-amplified and assembled separately onto histone octamers to reconstitute NCPs and evaluated by EMSA to ensure that in later rounds of selection (Round 2-4) the residual free DNA in the samples was also minimal (left lanes). To further test for the correctness of NCP reconstitution, the samples were digested by MNase, which indeed indicated full protection of their DNA (right lanes). Images are representative of the gels obtained in different rounds of PIONEAR-seq and using different TFs.

**d)** Evaluation of the effect of TF motif insertion on nucleosome positioning. To examine if the insertion of TF motifs affect the nucleosome positioning sequences, *NRCAM* templates, eventually carrying a CEBPB-specific motif (TTGCGCAA) at the indicated locations (red underlying segment) (**Supplementary Table 3**) were separately assembled into NCPs. After digestion with MNase (same reaction conditions as in (c)), DNA fragments were collected from protected NCPs, sequenced and mapped to the template to identify end cuts.

Distributions of absolute cut counts at the 5' (left) and 3' (right) end of the MNase-digested fragments are depicted. These distributions show that the insertion of a TF binding site, which modifies the distribution within the 8-bp of the insertion (red vs. gray underlying segments), does not modify the overall accessibility of the NCPs. Two independent replicates indicated that this evaluation is very reproducible. Images are representative of profiles obtained with the same assay on other templates.

**(e)** Evaluation of $N_{20}$ tile location in nucleosome positioning for PIONEAR-seq NCP input libraries. To examine whether the location of the random $N_{20}$ tile affects nucleosome positioning and accessibility, NCP input libraries were digested with MNase (same reaction conditions as in (c)). The DNA fragments from protected NCPs were collected, sequenced, selected to come from the *NRCAM*-based library, stratified according to the location of the random $N_{20}$ tile (blue) and analyzed for their end distributions. For each $N_{20}$ location, the relative distributions (%) of the cuts in the MNase-digested fragments at the left end and the right side of the random tile are shown (cut percentage color-coded as according to the legends for "Cut frequency (%)"). These distributions, which are very similar upon changes in the $N_{20}$ location, indicate that the insertion of the $N_{20}$ does not affect the overall accessibility profile of the NCPs. Two independent replicates indicated that this evaluation is very reproducible.

**(f)** For the indicated templates, same evaluation as in (e), representative of two independent replicates.

**Extended Data Fig. 2: Evaluating binding specificity to nucleosome DNA across a panel of TFs by PIONEAR-seq.**

**(a)** Anti-GST Western immunoblots were conducted to assay the production and the quality of the indicated GST-tagged full-length human TFs by using the prokaryotic *in vitro* transcription and translation PURExpress (NEB). The stars indicate the correct bands, corresponding to the full-length TFs.

**(b)** Comparison of the highest *k*-mer enrichment in libraries based on distinct templates (symbols), selected by the indicated TFs (or by the GST mock control) and assembled on NCPs (purple) versus kept as free DNA (yellow). Similar to Fig. 1b.

**(c, d)** Similar comparisons as in (b), but using the library's information gain quantified by the Kullback-Leibler instead of the highest *k*-mer enrichment. The GST threshold (equal to 0.04) was picked according to the highest value of the GST libraries to highlight TF libraries that exhibit an information gain due to TF specificity for DNA recognition.

(**e-h**) Systematic comparison between PIONEAR-seq and lig200 NCAP–SELEX data for binding specificity to nucleosome DNA. In published analyses of lig200 NCAP-SELEX data[18], the E-MI (enriched 3-mer pair based mutual information) diagonal signal is used to locate TF-DNA contacts without predefined assumptions on its binding specificity. The lig200 library contains a 154-bp random region in which the central 94 bp are expected to contact the nucleosome in the NCAP-SELEX assay. Therefore, the length of the central minimum of the E-MI diagonal signal reflects the portion of the random region where the nucleosome limits the TF-DNA interaction (*DNA region protected by the nucleosome* ). The shown E-MI diagonals are normalized so that the cumulative sum of each profile is equal to 1. Close paralogs found by using TFclass[32]. The DNA regions protected by the nucleosome from the invasion of TFs are depicted and quantified (red lines).

(**e**) E-MI diagonal signals for TFs (or close paralogs, as indicated) that recognize nucleosome DNA according to PIONEAR-seq (**Fig. 1b, Extended Data Fig. 2c**). For OCT4, which is a member of the POU5 subfamily[33], no subfamily members were assessed by lig200 NCAP–SELEX in Zhu *et al., Nature*, 2018.

(**f**) E-MI diagonal signals for TFs (or close paralogs) that do not recognize nucleosome DNA according to PIONEAR-seq (b, d). Note that for PU.1, which is a member of the ETS-III subfamily, and ZNF143, which is a member of the ZNF76-like family, no subfamily members were assessed by lig200 NCAP–SELEX[18].

(**g**) E-MI diagonal signals for two TFs previously highlighted[18] to characterize either binding specifically to nucleosomal DNA (VSX1) or exclusion from nucleosomal DNA in the presence of free DNA (SREBF2).

(**h**) Barplot of values of nucleosome protection from the invasion of the indicated TFs (as depicted in red in **(e-g)**). TFs from (e) exhibit similar values to the nucleosome DNA binder VSX1, while TFs from (b) exhibit similar values to the free DNA binder SREBF2.

**Extended Data Fig. 3: TFs recognize similar DNA motifs on free DNA as on nucleosomes.**

**(a)** For the indicated TFs, the correlation between the *k*-mer enrichment in W601 libraries selected either as free DNA (x-axis) or NCPs (y-axis) is quantified by the Spearman correlation coefficient $R_S$. The free DNA library came from earlier selection Rounds (1 to 3) so that their enrichment levels are more similar to those of NCP libraries from Round 4 selection for the same TF.

**(b, c)** 8-mer enrichment in free DNA (x-axis) versus NCPs (y-axis) libraries for the indicated libraries selected by (b) PAX7 and (c) RUNX2. Similar to (a) and Fig. 2a. Plots are representative of the 8-mer enrichment in the other templates used in PIONEAR-Seq. The most enriched 8-mers for each TF are highlighted and match the 8-mers used in (d).

**(d)** CEBPB, PAX7 and RUNX2 recognize their specific binding sites homogeneously along the random regions in the free DNA libraries. Positional density profiles of the indicated 8-mers in the free DNA libraries after 4 rounds of selection by the indicated TFs. Similar to Fig. 3a, but without the assembly of the DNA libraries onto NCPs.

**Extended Data Fig. 4: Predicted structures for CEBPB binding on W601-based NCPs by AlphaFold3**

AlphaFold3-predicted structures of the DNA binding domains of a CEBPB dimer binding to its cognate 8-mer inserted between the indicated base pairs of the W601 nucleosome. Top row is the same as Fig. 2c. On the right, the Predicted aligned error (PAE) plots associated with each predicted structure produced by AlphaFold3. PAE plots measure how confident AlphaFold2 is in the relative position of two residues within the predicted structure. Domains that are well packed and have a correct relative placement in the predicted structure have lower Expected Position Error (color bar).

**Extended Data Fig. 5: EMSAs for the interaction of W601-based NCPs with CEBPB**

Representative EMSA gels showing CEBPB binding to W601 sequences, either as free DNA (middle gels) or NCPs (top and bottom gels), carrying CEBPB-specific binding motifs at the indicated positions. CEBPB was purified either as DBD (top and middle gels) or as full-length protein (bottom gel). Similar EMSA reaction conditions as in Fig. 2e.

**Extended Data Fig. 6: Cyclizability of nucleosomal template sequences regulates TF binding to specific nucleosome ends.**

**(a)** Percentages of sequences from each of the W601*, NRCAM, ALBN1, CX3CR1* NCP libraries before and after MNase digestion and sequencing (two replicates each).

**(b)** Sequence cyclizability profiles of the W601*, NRCAM, ALBN1, CX3CR1* template predicted by the tool DNAcycP[34].

**(c)** Averaged cyclizability of sequences with random tiles at different positions in each of the W601, *NRCAM, ALBN1, CX3CR1* libraries.

**(d-f)** EMSAs for the interaction of *NRCAM*-based NCPs with CEBPB. Representative EMSAs showing CEBPB binding to W601 sequences, either as NCPs (d, e) or free DNA (f), carrying CEBPB-specific binding motifs at the indicated positions. CEBPB was either purified as DBD (d, f) or as full-length (e). Similar EMSA reaction conditions as in Fig. 3f.

**Extended Data Fig. 7: Positional density profiles of TF binding for OCT4, FOXA1, GATA1 and GATA4.**

For the indicated TFs, a visual comparison of the positional density profiles in NCP (top) vs. free DNA (bottom) libraries after 4 Rounds of TF selection. The resulting, most enriched *k*-mers are shown for the TFs.

**Extended Data Fig. 8: Comparison of positional binding preference called by PIONEAR-seq vs. NCAP-SELEX**

**(a)** For the indicated TFs, the positional density profiles of the top two enriched *k*-mers in W601-based PIONEAR-seq libraries after 4 Rounds of TF selection (same profiles as in **Fig. 3a** and **Extended Data Fig. 7**).

**(b)** For the TFs (or close paralogs) shown in (a), the published E-MI diagonal profiles of lig147 libraries selected by NCAP-SELEX (*i.e.*, NCPs) or HT-SELEX (free DNA)[18]. The lig147 NCAP-SELEX data were used previously to call non-mutually exclusive types of positional preferences[18], here end, dyad and periodic binding (labeled in red).

**Extended Data Fig. 9: DNA cyclizability of the sequences surrounding TF binding sites influences TF binding to nucleosomes.**

**(a)** Schema of analyzing lig147 TF-SELEX library. Input DNA ligands of 147-bp were bound by nucleosomes, TFs, or nucleosomes and TFs, respectively for the Nucleosome-SELEX library, TF HT-SELEX library, and TF NCAP-SELEX library. Enriched 7-mers were discovered *de novo* from TF NCAP-SELEX libraries and used to filter the libraries. The internal region of the sequence was defined as the region without overlap with the primers and containing the TF-specific binding motif.

**(b)** Sequence cyclizability of end-binding reads in the CEBPB NCAP-SELEX library with internal cyclizability fitted by linear regression. The internal cyclizability slope is defined as the slope of the fitted linear regression.

**(c)** TF family distribution of end binders (left) and non-end binders (right).

**(d)** Linear regression slopes of the internal sequence cyclizability and the average sequence cyclizability of end binding sequences from the Nucleosome-SELEX, NCAP-SELEX, and HT-SELEX libraries for end binding TFs (left) and non-end binding TFs (right).

**(e)** Average periodicity intensity of nucleotide ‘A’ in two symmetrical regions on the nucleosomes across filtered reads from the Nucleosome-SELEX library, the TF NCAP-SELEX library, and the TF HT-SELEX library, for end-binding (left) and non-end-binding TFs (right).

(**f**) Averaged profiles of sequence cyclizability in CEBPB NCAP-SELEX data, oriented and stratified to show end binding of specific CEBPB 8-mers at progressively more internal regions from the nucleosome ends (25-35, 30-40, 35-45, and 40-50 bp, as color coded).

(**g**) For each end binder (**Supplementary Table 8**), the average profile was calculated as in (f). All the profiles for the same distances of the binding *k*-mers from the nucleosome ends were averaged and displayed.

### 4 - Supplementary References


1 Dyer, P. N. *et al.* Reconstitution of nucleosome core particles from recombinant histones and DNA. *Methods Enzymol* **375**, 23-44, doi:10.1016/s0076-6879(03)75002-2 (2004).

2 Fernandez Garcia, M. *et al.* Structural Features of Transcription Factors Associating with Nucleosome Binding. *Mol Cell* **75**, 921-932 e926, doi:10.1016/j.molcel.2019.06.009 (2019).

3 Lowary, P. T. & Widom, J. New DNA sequence rules for high affinity binding to histone octamer and sequence-directed nucleosome positioning. *J Mol Biol* **276**, 19-42, doi:10.1006/jmbi.1997.1494 (1998).

4 Zhang, J., Kobert, K., Flouri, T. & Stamatakis, A. PEAR: a fast and accurate Illumina Paired-End reAd mergeR. *Bioinformatics* **30**, 614-620, doi:10.1093/bioinformatics/btt593 (2014).

5 Berger, M. F. *et al.* Compact, universal DNA microarrays to comprehensively determine transcription-factor binding site specificities. *Nat Biotechnol* **24**, 1429-1435, doi:10.1038/nbt1246 (2006).

6 Riley, T. R. *et al.* SELEX-seq: a method for characterizing the complete repertoire of binding site preferences for transcription factor complexes. *Methods Mol Biol* **1196**, 255-278, doi:10.1007/978-1-4939-1242-1_16 (2014).

7 Rastogi, C., Liu, D., Melo, L. & Bussemaker, H. J. SELEX: Functions for analyzing SELEX-seq data. doi:10.18129/B9.bioc.SELEX, R package version 1.34.0, https://bioconductor.org/packages/SELEX. (2023).

8 Andersen, K. R., Leksa, N. C. & Schwartz, T. U. Optimized E. coli expression strain LOBSTR eliminates common contaminants from His-tag purification. *Proteins* **81**, 1857-1861, doi:10.1002/prot.24364 (2013).

9 Abramson, J. *et al.* Accurate structure prediction of biomolecular interactions with AlphaFold 3. *Nature* **630**, 493-500, doi:10.1038/s41586-024-07487-w (2024).

10 Wang, H., Xiong, L. & Cramer, P. Structures and implications of TBP-nucleosome complexes. *Proc Natl Acad Sci U S A* **118**, doi:10.1073/pnas.2108859118 (2021).

11 Chen, S., Lei, M., Liu, K. & Min, J. Structural basis for specific DNA sequence recognition by the transcription factor NFIL3. *J Biol Chem* **300**, 105776, doi:10.1016/j.jbc.2024.105776 (2024).

12 Luger, K., Mader, A. W., Richmond, R. K., Sargent, D. F. & Richmond, T. J. Crystal structure of the nucleosome core particle at 2.8 A resolution. *Nature* **389**, 251-260, doi:10.1038/38444 (1997).

13 Li, S., Olson, W. K. & Lu, X. J. Web 3DNA 2.0 for the analysis, visualization, and modeling of 3D nucleic acid structures. *Nucleic Acids Res* **47**, W26-W34, doi:10.1093/nar/gkz394 (2019).

14 Yang, J. *et al.* Structural basis for effects of CpA modifications on C/EBPbeta binding of DNA. *Nucleic Acids Res* **47**, 1774-1785, doi:10.1093/nar/gky1264 (2019).

15 Basu, A. *et al.* Measuring DNA mechanics on the genome scale. *Nature* **589**, 462-467, doi:10.1038/s41586-020-03052-3 (2021).

16 Park, J. & Ha, T. J. In preparation. (2024).

17 Li, K., Carroll, M., Vafabakhsh, R., Wang, X. A. & Wang, J. P. DNAcycP: a deep learning tool for DNA cyclizability prediction. *Nucleic Acids Res* **50**, 3142-3154, doi:10.1093/nar/gkac162 (2022).

18 Zhu, F. *et al.* The interaction landscape between transcription factors and the nucleosome. *Nature* **562**, 76-81, doi:10.1038/s41586-018-0549-5 (2018).

19 Luo, Y. *et al.* New developments on the Encyclopedia of DNA Elements (ENCODE) data portal. *Nucleic Acids Res* **48**, D882-D889, doi:10.1093/nar/gkz1062 (2020).

20 Chen, K. *et al.* DANPOS: dynamic analysis of nucleosome position and occupancy by sequencing. *Genome Res* **23**, 341-351, doi:10.1101/gr.142067.112 (2013).

21 Martins, A. L., Walavalkar, N. M., Anderson, W. D., Zang, C. & Guertin, M. J. Universal correction of enzymatic sequence bias reveals molecular signatures of protein/DNA interactions. *Nucleic Acids Res* **46**, e9, doi:10.1093/nar/gkx1053 (2018).
22 Hinrichs, A. S. *et al.* The UCSC Genome Browser Database: update 2006. *Nucleic Acids Res* **34**, D590-598, doi:10.1093/nar/gkj144 (2006).
23 Heinz, S. *et al.* Simple combinations of lineage-determining transcription factors prime cis-regulatory elements required for macrophage and B cell identities. *Mol Cell* **38**, 576-589, doi:10.1016/j.molcel.2010.05.004 (2010).
24 Quinlan, A. R. & Hall, I. M. BEDTools: a flexible suite of utilities for comparing genomic features. *Bioinformatics* **26**, 841-842, doi:10.1093/bioinformatics/btq033 (2010).
25 Zhou, Q. *et al.* ZNF143 mediates CTCF-bound promoter-enhancer loops required for murine hematopoietic stem and progenitor cell function. *Nat Commun* **12**, 43, doi:10.1038/s41467-020-20282-1 (2021).
26 Magnitov, M. D. *et al.* ZNF143 is a transcriptional regulator of nuclear-encoded mitochondrial genes that acts independently of looping and CTCF. *biorxiv*, doi:https://doi.org/10.1101/2024.03.08.583864 (2024).
27 Soufi, A. *et al.* Pioneer transcription factors target partial DNA motifs on nucleosomes to initiate reprogramming. *Cell* **161**, 555-568, doi:10.1016/j.cell.2015.03.017 (2015).
28 Minderjahn, J. *et al.* Mechanisms governing the pioneering and redistribution capabilities of the non-classical pioneer PU.1. *Nat Commun* **11**, 402, doi:10.1038/s41467-019-13960-2 (2020).
29 Lian, T., Guan, R., Zhou, B. R. & Bai, Y. Structural mechanism of synergistic targeting of the CX3CR1 nucleosome by PU.1 and C/EBPalpha. *Nat Struct Mol Biol* **31**, 633-643, doi:10.1038/s41594-023-01189-z (2024).
30 Michael, A. K. *et al.* Mechanisms of OCT4-SOX2 motif readout on nucleosomes. *Science* **368**, 1460-1465, doi:10.1126/science.abb0074 (2020).
31 Yu, X. & Buck, M. J. Defining TP53 pioneering capabilities with competitive nucleosome binding assays. *Genome Res* **29**, 107-115, doi:10.1101/gr.234104.117 (2019).
32 Wingender, E., Schoeps, T., Haubrock, M., Krull, M. & Donitz, J. TFClass: expanding the classification of human transcription factors to their mammalian orthologs. *Nucleic Acids Res* **46**, D343-D347, doi:10.1093/nar/gkx987 (2018).
33 Malik, V., Zimmer, D. & Jauch, R. Diversity among POU transcription factors in chromatin recognition and cell fate reprogramming. *Cell Mol Life Sci* **75**, 1587-1612, doi:10.1007/s00018-018-2748-5 (2018).
34 Li, G. & Widom, J. Nucleosomes facilitate their own invasion. *Nat Struct Mol Biol* **11**, 763-769, doi:10.1038/nsmb801 (2004).
